# Regulatory co-option of a homeobox gene drives parasitoid venom evolution

**DOI:** 10.64898/2026.06.22.732516

**Authors:** Yi Yang, Shasha Wang, Chang Liu, Deqing Yang, Shan Xiao, Zhichao Cao, Shuxing Lao, Yaoyao Chen, Qi Fang, Gongyin Ye, Xinhai Ye

## Abstract

How gene regulatory networks are rewired to generate phenotypic and functional innovation remains a central question in evolutionary biology. Parasitoid wasp venoms provide a powerful system for addressing this question, as their repertoires evolve rapidly through extensive lineage-specific turnover, yet the regulatory principles underlying such flexibility are largely unknown. Here we integrate tissue-resolved transcriptomic, chromatin-accessibility and histone-modification profiling to reconstruct the venom regulatory network of the parasitoid wasp *Pteromalus puparum*. We show that venom expression is embedded in distinct chromatin states and shaped by regulatory elements associated with venom-gland transcription. Comparative and functional analyses support a general model in which regulators related to the endoplasmic reticulum stress and unfolded protein response pathways have been repeatedly recruited to venom regulation across venomous lineages. Unexpectedly, we identify the recently co-opted homeobox gene *Lbx* as a lineage-specific hub that regulates more than half of venom genes and is linked to enhancer evolution. These results reveal a nested model of venom regulatory evolution, in which an ancestral secretory programme provides a reusable regulatory backbone, while newly co-opted homeobox gene specify a lineage-specific venom expression. Our study highlights regulatory co-option as a mechanism by which conserved developmental genes can acquire new physiological functions during adaptive evolution.

## Main

Evolutionary innovation in organ phenotype and function often arises through the emergence and rewiring of gene regulatory networks (GRNs)^1–3^. Such innovation frequently involves the co-option of pre-existing regulators together with the reorganization of their downstream networks, thereby generating novo functions without requiring the entirely new regulatory logic^4–8^. One of the clearest examples is homeobox genes, which, although ancestrally central to body patterning and organogenesis^9,10^, have been reported to be recruited beyond their conserved functions and rewired into novel regulatory programmes to generate novel phenotypic innovations across diverse animal linages, for example in butterfly eyespots evolution^6,11,12^, marsupial gliding membranes evolution^13^, and fin to limb transition in vertebrate evolution^14^. However, how such regulators are incorporated into organ-specific GRNs to drive organ function innovation is largely unknown.

Venom systems offer a compelling setting to address this problem^15^. Venoms have originated independently more than 100 times across the animal tree of life^15,16^, yet each case requires GRN innovation to generate lineage-specific venom repertoires for different purpose, such as prey in snakes^17^ and reproduction in parasitoid wasps^18^. Parasitoid wasps are especially informative in this case because their venoms evolve rapidly even over short evolutionary timescales^19–21^. For instance, in two *Anastatus* wasps that diverged only ∼3 million years ago, more than half of venom genes show difference^22^, highlighting the evolutionary lability of venom gene repositories despite conservation of the venom gland itself. This combination of organ conservation and rapid functional divergence makes parasitoid venoms an excellent system for dissecting how organ-specific regulatory programmes evolve.

Despite extensive work on venom composition, function, and evolution, the regulatory architecture underlying venom gene expression and how venom GRN evolve to regulate gene expression remain largely unresolved^15,21^. Work in snakes has begun to identify candidate trans-regulatory factors associated with venom expression, including ATF6, XBP1, GRHL1, and EHF, and has suggested that venom GRNs were assembled in part through the co-option of ancient pathways, particularly the ER stress and unfolded protein response (UPR) signaling pathways^23–26^. These findings, together with comparative transcriptomic analyses of venom glands, suggest that venomous animals share a conserved regulatory programme, likely inherited from the ancestral secretory cells^27,28^. However, it remains unclear whether a similar regulatory framework exists in parasitoid wasps, and what additional mechanisms might drive the evolution of venom GRNs in this group.

In this study, we construct a multi-omic atlas of transcriptional and epigenomic regulation across the venom gland and four non-venom tissues of the parasitoid wasp *P. puparum*, to resolve the regulatory architecture of its venom system. We show that venom genes are expressed in a highly tissue-specific manner and are coupled to distinct chromatin states and histone modifications. By integrating transcriptomic and epigenomic information, we reconstruct a venom GRN that implicates regulators related to the ER stress and UPR response, consistent with a conserved secretory backbone also observed in other venomous lineages. We further uncover the recent co-option of the homeobox gene *Lbx* as a central regulator of venom expression, likely driven by *cis-*regulatory innovation in an enhancer. These findings support a nested model of venom GRN evolution, in which an ancestral secretory related framework provides a foundation, and the co-option and rewiring of deeply conserved developmental regulators to enable venom function innovations. Our results also highlight the post-development expression and non-development related function of homeobox gene, providing new insights to the functional diversification of homeobox gene.

## Results

### An atlas of tissue-specific transcriptional and epigenomic landscapes in a parasitoid wasp

To comprehensively characterize the transcriptional and epigenomic landscapes of *P. puparum*, we generated 12 new RNA-seq libraries (extending our previous venom gland data^29^), together with 15 ATAC-seq (Assay for Transposase-Accessible Chromatin using sequencing) and 30 CUT&Tag (Cleavage Under Targets and Tagmentation) libraries for H3K4me3 and H3K27ac, across five female adult tissues: head, thorax, digestive tract, ovary, and venom gland of female adults (Fig. 1a; Supplementary Table 1-3). All epigenomic datasets exhibited high signal-to-noise ratios, with strong enrichment around transcription start sites (TSS) and an average FRiP (Fraction of reads in peaks) score of 0.42 (Supplementary Fig. 1a-d). Principal component analysis and clustering further separated samples by tissue type and showed high concordance among biological replicates (Supplementary Fig. 1e,f). These results together indicate the high quality and reproducibility of our multi-omics datasets.

**Figure 1.**
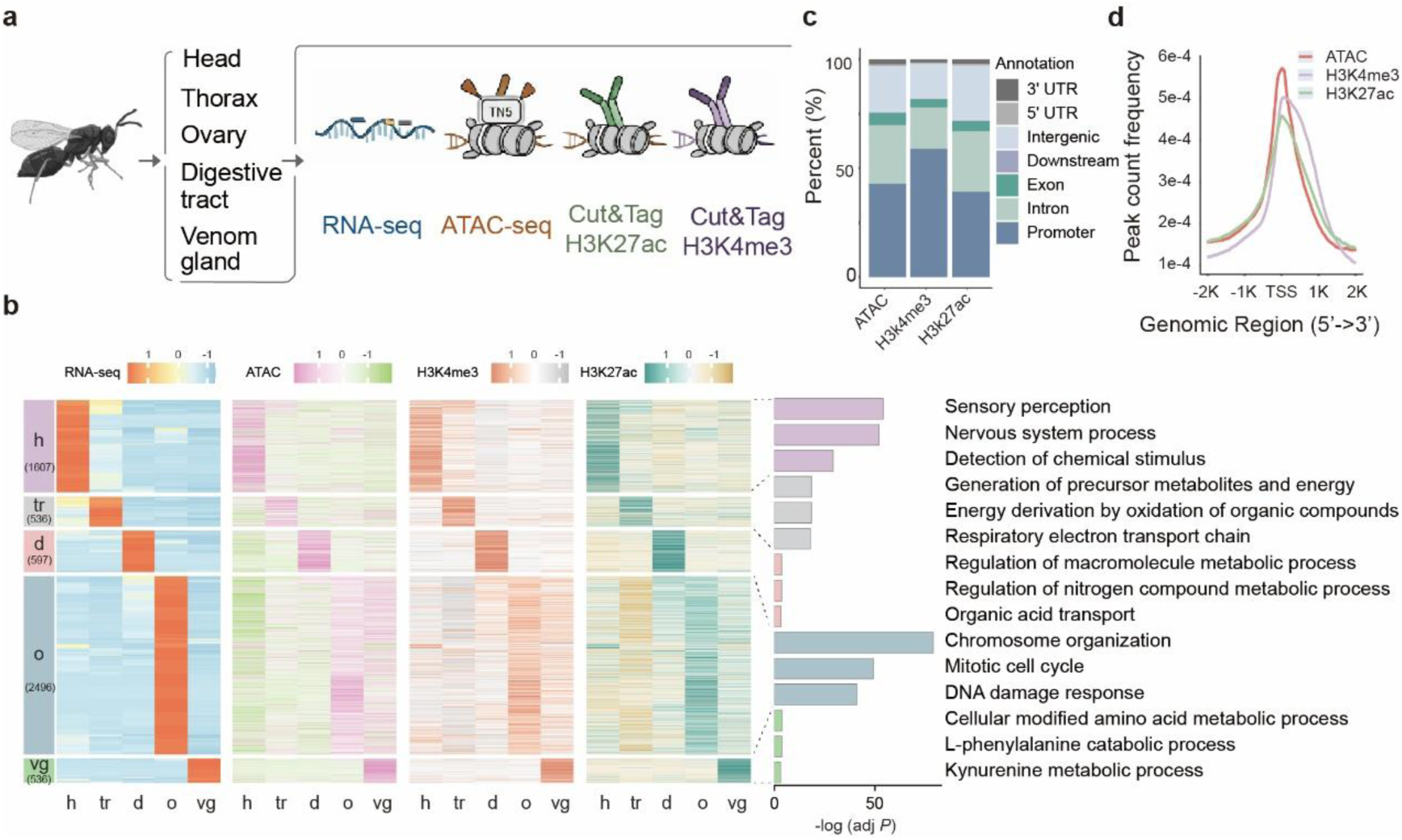
Transcriptional and epigenomic landscapes across tissues. (a) Schematic overview of the sampled tissues and multi-omics datasets generated in this study, including RNA-seq, ATAC-seq, and CUT&Tag profiling for H3K27ac and H3K4me3. (b) Heatmaps showing transcriptome, chromatin accessibility, H3K27ac, and H3K4me3 patterns for tissue-specific genes across he five tissues: head (h), thorax (tr), digestive tract (d), ovary (o), and venom gland (vg). RNA-seq data are represented as *Z*-scores of log₂(TPM + 1), while ATAC-seq and CUT&Tag data are represented as *Z*-scores of log₂(CPM + 1). Right-hand panels show Gene Ontology (GO) enrichment for tissue-specific genes. (c) Genome-wide annotation of accessible chromatin (ATAC) and histone modifications (H3K4me3, H3K27ac), showing their distribution across genomic features. (d) Aggregate profiles of ATAC, H3K4me3, and H3K27ac peaks around transcription start sites (TSS).

We detected 12,251 genes expressed (TPM > 1) in at least one tissue, representing 66% of all annotated genes in the *P. puparum* reference genome^29,30^. Among these, 9,103 genes (∼50% of annotated) exhibited differential expression in at least one tissue, whereas 3,255 genes were consistently expressed across all tissues (Extended Data Fig. 1a). We further used the Tau metric to define genes with tissue-specific expression and identified 5,739 tissue-specific genes across the five tissues (Fig. 1b). Gene ontology (GO) enrichment analysis showed that tissue-specific genes are predominantly associated with functions relevant to their respective tissues, whereas consistently expressed genes are enriched in processes linked to fundamental cellular functions (Fig. 1b, Extended Data Fig. 1b). For instance, the 1,607 head-specific genes are primarily involved in sensory perception and nervous system processes, whereas the 2,496 ovary-specific genes are enriched in mitotic cell cycle and chromosome organization processes (Fig. 1b).

We next profiled the chromatin states and histone modifications in the *P. puparum* genome. Specifically, we identified 62,441 ATAC peaks, 19,779 H3K4me3 peaks, and 38,801 H3K27ac peaks across the five tissues, collectively covering ∼17.3% of the genome (Extended Data Fig. 1c). These peaks exhibited conserved distribution patterns similar to those in other animals, with the majority located in promoter regions. However, ATAC and H3K27ac peaks were more prominently situated in intronic and intergenic regions (Fig. 1c,d). This distribution pattern aligns with their known functions of these chromatin features: H3K4me3 histone modifications are typically associated with active promoters, while H3K27ac marks and ATAC are markers in both promoter and enhancer regions in animals^31,32^. We then compared the peaks across tissues and identified tissue-specific peaks, and found that genes associated with these peaks generally enriched for tissue-related functions (Extended Data Fig. 1d; Supplementary Fig. 2).

Given that chromatin states and histone modifications play critical roles in regulating gene expression^31^, we next explored the relationship between gene expression dynamics and chromatin stats. We first identified genes associated with ATAC, H3K4me3, and H3K27ac peaks, respectively.

Genes with these chromatin features exhibited significantly higher expression levels in their corresponding tissues (Extended Data Fig. 1e, *P* < 0.0001, one-side Mann-Whitney *U* test). Furthermore, we calculated the Pearson correlation coefficients (PCC) between expression levels and the signal intensity of each chromatin feature for tissue-specific genes. We observed strong positive correlations between chromatin accessibility, H3K4me3, H3K27ac, and gene expression levels (Extended Data Fig. 1f). These results are consistent with the reported functions of these chromatin stats in activating gene expression^31^, suggesting that gene expression in *P. puparum* is coordinated by chromatin states and histone modifications.

### Venom gene expression is associated with open chromatin and active histone modifications

We previously integrated multi-omics data (PacBio Iso-Seq, CAGE-seq, and proteomics) to identify 90 high-confidence venom protein coding genes (hereafter referred to as venom genes) with precise TSS information in *P. puparum*^29^. Using the transcriptomic and epigenomic datasets generated in this study, we further profiled the regulatory landscapes of these venom genes. Our analysis revealed that most venom genes (76/90, 84%) are highly expressed in the venom gland relative to other tissues (Fig. 2a-c; log2FC > 1, adjusted *P* value < 0.05; Supplementary Table 4). Among these, 61 venom genes exhibited a Tau > 0.8, indicating venom-gland-specific expression (Fig. 2b,C; Supplementary Table 4). We then examined chromatin accessibility and active histone modifications at venom gene loci. The majority of venom genes (73 of 90, 81%) harboured at least one ATAC peak together with H3K4me3 and H3K27ac peaks within their promoter or genic regions (Fig. 2c; Supplementary Table 5).

**Figure 2.**
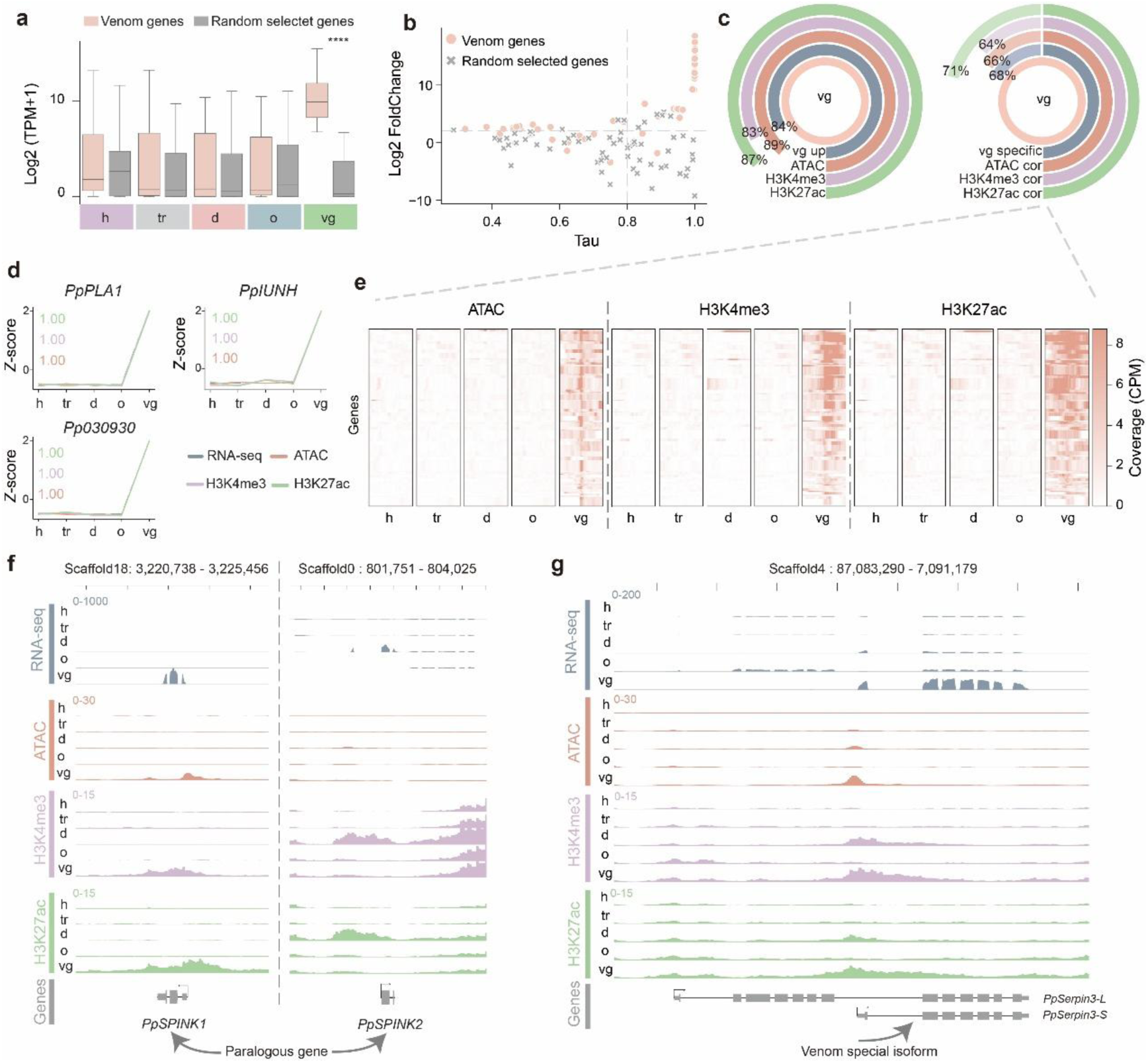
Coordinated transcriptional and epigenomic activation of venom genes in the venom gland. (a) Expression of venom genes (red) and randomly selected control genes (grey) across tissues. **** *P* < 0.0001, one-sided Mann-Whitney *U* test, n =90 genes per group. (b) Tissue specificity and venom-gland biased expression of venom genes. The tissue-specificity index Tau is plotted against the log2 fold change in venom gland relative to non-venom tissues for venom genes (red) and control genes (grey). (c) Proportions of venom-gland-upregulated genes (left, vgup) and venom-gland-specific genes (right, vgss) associated with chromatin accessibility and active histone modifications. Left, fractions of genes overlapping ATAC-seq, H3K4me3 and H3K27ac peaks. Right, fractions of venom genes whose expression is positively correlated with ATAC-seq, H3K4me3 and H3K27ac signals (PCC > 0.7). (d) Representative venom genes showing coordinated venom-gland-specific expression, chromatin accessibility and active histone marks across tissues. (e) Heatmaps of ATAC-seq, H3K4me3 and H3K27ac signal across venom specific genes in the indicated tissues, highlighting enrichment in venom gland. (f) Genome-browser views of representative paralogous loci (left) and an example of isoform-level co-option of a venom gene (right), illustrating venom-gland-specific gene activation accompanied by increased chromatin accessibility and active histone modifications.

To determine whether these chromatin features covary with venom-gland-biased transcription, we calculated PCCs between the expression profile of each venom gene and its corresponding ATAC, H3K4me3 and H3K27ac signals. We found a coordinated pattern between venom-gland expression and active chromatin states, with 66%, 64% and 71% venom genes showing strong positive correlations (PCC > 0.7; Fig. 2c-e; Supplementary Table 5). Venom genes are reported to origin through co-option, we next compared the chromatin profiles of venom genes and its non-venom paralogues and found correlations. For example, *PpSPINK1* (OGS id: *Pp132210*) and *PpSPINK2* (OGS ID: *Pp139100*) are paralogous genes; *PpSPINK1* is specifically expressed in the venom gland, whereas *PpSPINK2* is predominantly expressed in the digestive tract. Notably, both genes exhibited concordant ATAC-seq and histone modification signals (H3K4me3 and H3K27ac) at their respective TSS regions (Fig. 2f). Moreover, co-option of venom genes has been reported at the isoform level, and we previously demonstrated that parasitoid wasps specifically co-opt a short serpin isoform (*Ppserpin-S*) for venom function^21,29^. Our epigenomic data align well with this observation, as we detected significant ATAC and H3K4me3/H3K27ac signals around the TSS of *Ppserpin-S*, but not around that of the long isoform *(Ppserpin-L*, Fig. 2g). Collectively, these results provide evidence that venom genes are specifically and highly expressed in the venom gland, and that this expression pattern is associated with permissive chromatin states and activating histone modifications.

### Rapid sequence turnover of venom-associated *cis-*regulatory elements

Previous studies have shown that parasitoid wasp venoms undergo rapid evolutionary turnover, whereas signatures of selection are rarely detected in protein coding sequences^19,22^. We therefore asked whether this pattern is accompanied by rapid evolution of *cis*-regulatory elements. To identify *cis-*regulatory elements in the venom gland of *P. puparum*, we integrated chromatin accessibility data with H3K4me3 and H3K27ac profiles generated in this study. Consistent with the conserved functions and distribution patterns of histone modigications^31^, we observed significant co-occurrence of ATAC-seq signals with both H3K4me3 (*Z*-score: 320, *P* < 0.0001; Permutation test) and H3K27ac signals (*Z*-score: 296, *P* < 0.0001; Permutation test; Supplementary Fig. 3). Accordingly, we assigned ATAC peaks co-localized with both H3K4me3 and H3K27ac signals as putative active promoters (pAPs), and those overlapping with H3K27ac but lacking H3K4me3 were designated as putative active enhancers (pAEs) (Fig. 3a). In total, we identified 6,764 pAPs and 2,979 pAEs in the venom gland (Fig. 3b). The genomic distribution of these elements aligned with expected functional roles: the majority of pAPs (85%) were located in the promoter regions, whereas most pAEs (53%) were located within introns or intergenic regions (Fig. 3c). Among them, 96 pAPs and 71 pAEs were associated with venom genes (designated vpAPs and vpAEs, respectively), collectively covering 67.8% (61/90) and 37.8% (34/90) of venom gene (Supplementary Table 6).

**Figure 3.**
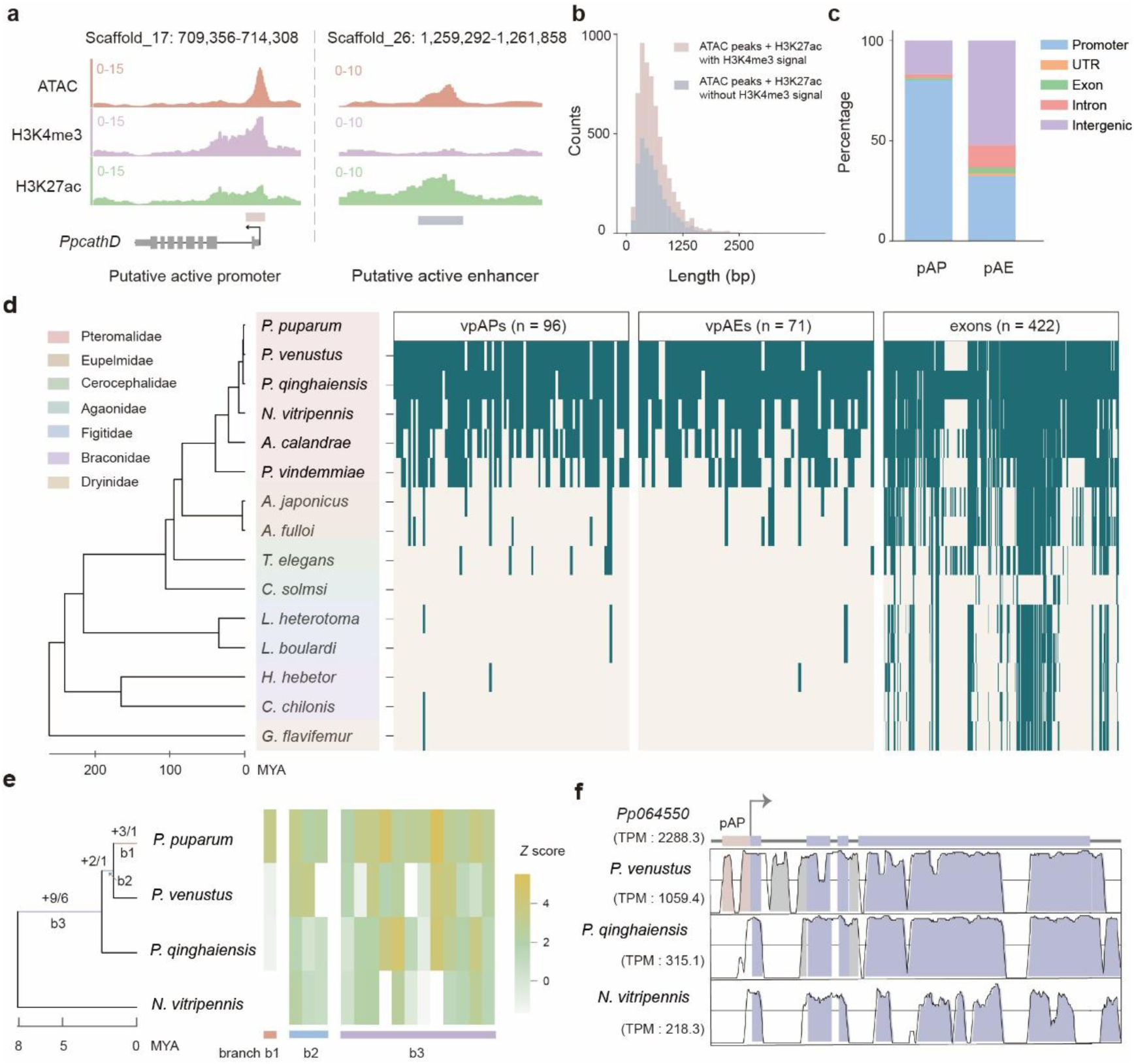
Identification and evolutionary dynamics of cis-regulatory elements associated with venom genes. (a) Representative loci illustrating the chromatin features used to define putative active promoters (pAPs) and putative active enhancers (pAEs). (b, c) Length distributions (b) and genomic annotations (c) of identified pAPs and pAEs. (d) Evolutionary analysis of the sequence conservation of pAPs, pAEs and exon of venom genes across parasitoid wasps. The species phylogenetic tree was adopt from previous studies^20^. The heatmaps indicate whether each element is alignable in each species. (e) Evolutionary turnover of vpAPs and vpAEs across four closely related parasitoid wasps. Numbers at internal nodes indicate branch-specific gains of vpAPs and vpAEs. Right, heatmap showing venom-gland expression of venom genes and their orthologues across species. (f) Representative locus showing a lineage-gained pAP and venom-gland expression of the associated gene and its orthologues across species.

To explore the evolutionary dynamics and potential regulatory roles of these elements in venom gene evolution, we generated 15-way whole-genome alignments among parasitoid wasps with available venom gland RNA-seq data (spanning 4 superfamilies and 7 families; Fig. 3d; Supplementary Table 7), and traced their sequence conservation across phylogenetically diverse taxa. Overall, vpAP and vpAE showed significantly lower alignability than coding regions across species (Fig 3d, Supplementary Fig. 4; one-sided Mann-Whitney *U* test, vpAPs vs. CDS: *P* = 1.46e- 14; vpAEs vs. CDS: *P* = 1.02e-12), suggesting rapid turnover of venom-associated regulatory elements. Notably, the alignability of both vpAPs and vpAEs declined dramatically beyond the six Pteromalidae species (*P. puparum* - *P. vindemmiae*), with only 11 pAPs (11.5%) and 8 pAEs (11.3%) traceable outside this clade (Fig. 3d; Supplementary Fig. 4).

Because most venom-associated *cis*-regulatory elements were detectable only within Pteromalidae, we next tested whether lineage-specific gains of these elements were associated with increased expression of nearby venom genes across four closely related Pteromalidae species that diverged approximately 8 million years ago (Fig. 3e). Overall, we found partial support for an association between lineage-specific element gains and expression divergence, with several loci exhibiting consistent patterns (Fig. 3e). For example, we identified an active promoter specific to *P. puparum* and *P. venustus*; its downstream gene, *Pp064550*, showed 6.28-fold higher expression than its orthologues in the other species (Fig. 3f). Together, our results indicate that venom-associated *cis*-regulatory elements undergo rapid sequence turnover, potentially playing a role in activating venom gene expression and thereby facilitating venom evolution in parasitoid wasps.

### Parallel evolution of venom associated regulators

We next asked how GRNs have been rewired in the venom systems of parasitoid wasps. While much is known about venom genes, including their composition, function, and origin^21^, our understanding of the regulators controlling their expression remains limited. To fill this gap, we identified 757 transcription factors (TFs) in the *P. puparum* genome (Supplementary Fig. 5) and used transcriptomic and epigenomic data to build five initial input GRNs, capturing complementary functional and physical evidence of regulation, including co-expression, GENIE3-inferred regulatory associations, chromatin states, TF-binding motifs, and ATAC-seq footprinting (Extended Data Fig. 2a). We integrated these networks using supervised machine learning and a scaled unsupervised framework, generating an integrative GRN (iGRN) comprising 718 TFs and 361,466 regulatory interactions (Extended Data Fig. 2a,b). The resulting iGRN outperformed each input layer and was supported by conserved *Drosophila* regulatory interactions as well as expression and functional coherence among TF co-regulated genes (Extended Data Fig. 2c-f), supporting its reliability and biological relevance.

We next interrogated the venom-associated subnetwork and identified 29 TFs with at least five predicted venom gene targets (Fig. 4a). Among these, 13 TFs showed elevated expression in the venom gland relative to non-venom tissues and were further supported by chromatin accessibility and histone modification profiles (Fig. 4b,c; Supplementary Table 8). To determine whether these candidate regulators exhibit conserved venom-gland activity across parasitoid wasps, we compared their expression profiles across lineages with sufficient sampling (i.e., ≥3 replicate transcriptome samples both in venom-gland and non-venom-gland replicates). In each species, 4 to 12 candidate TFs were significantly up-regulated in the venom gland. Among them, ATF6, CrebA, and GATA5 showed conserved venom-gland up-regulation across all parasitoid species examined, suggesting a shared regulatory logic underlying venom production in parasitoid wasps (Supplementary Fig. 6). We next ask whether venom gene regulation may show broader parallels across independently evolved venom systems. We compared the regulators identified in parasitoid wasps with those reported in snakes and found five overlapped candidate regulators (Fig. 4d). The most active regulators again included ATF6 and CrebA, which are core regulators related to ER stress and UPR response.

**Fig. 4.**
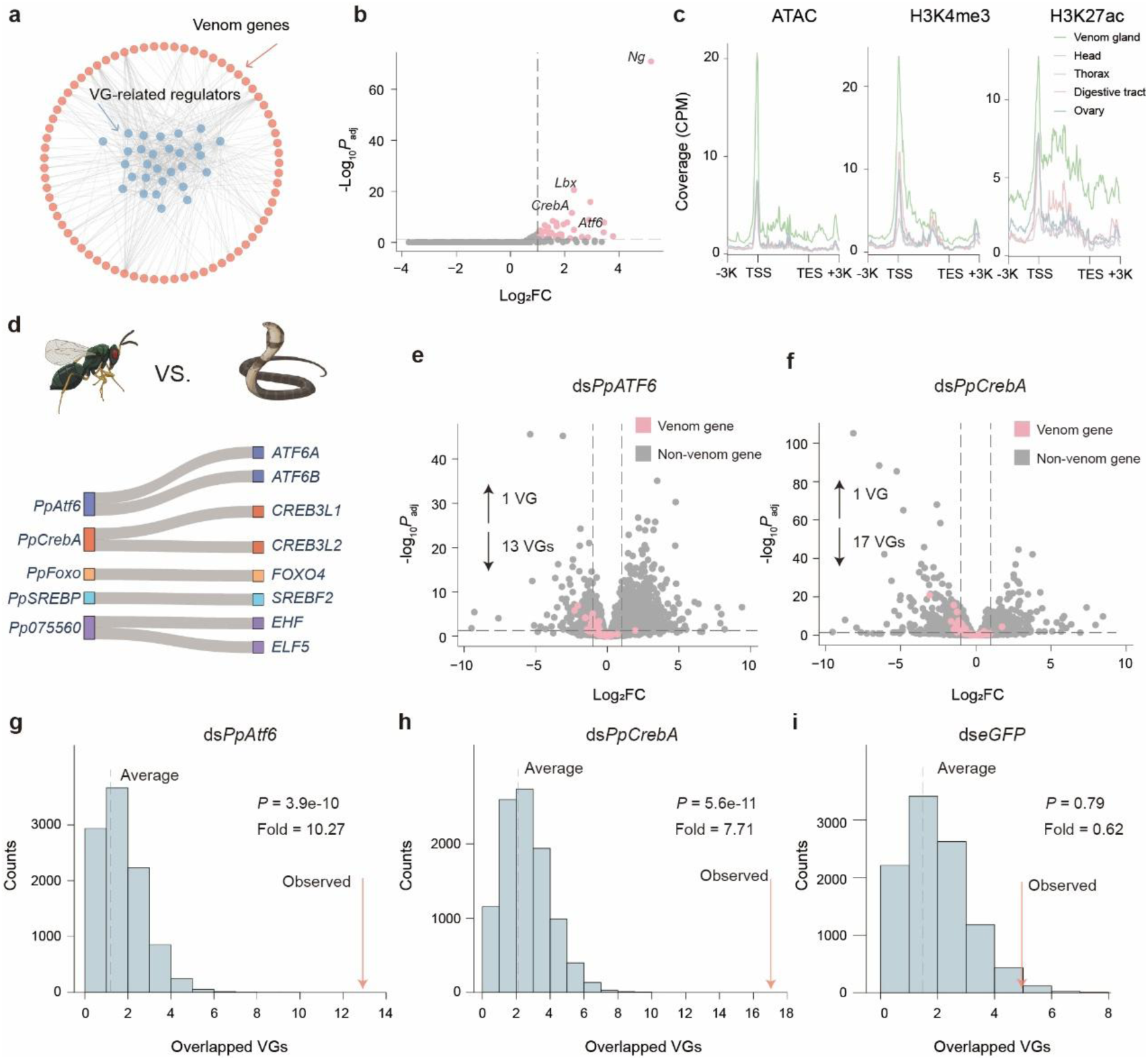
Venom gene regulatory network identifies candidate regulators and reveals parallels with snake venom regulation. (a) Venom-associated subnetwork derived from the iGRN, showing connections between venom genes (red) and related regulators (blue). (b) Volcano plot showing venom-gland-upregulated expression of candidate regulators. (c) Metagene profiles of ATAC-seq, H3K4me3 and H3K27ac signals across venom-related regulators in the venom gland. (d) Comparison of venom-related regulators identified in this study with regulators previously implicated in snake venom regulation. (e, f) Volcano plots showing transcriptomic changes after knockdown of *PpAtf6* (e) and *PpCrebA* (f), with venom genes highlighted in pink and non-venom genes in grey. (g-i) Permutation analyses comparing the observed number of downregulated venom genes with the null distribution expected from random sampling (n = 10,000) after knockdown of *PpAtf6* (g), *PpCrebA* (h) or *eGFP* (i). Arrows indicate the observed overlaps, and *P* values were calculated by permutation tests. The observed overlaps were greater than expected for *PpAtf6* and *PpCrebA* knockdown, but not for the *eGFP* control.

An important alternative explanation, however, is that these TFs reflect the general physiological demands of a highly secretory tissue, rather than specific involvement in venom gene regulation. To address this, we selected the two representative TFs, ATF6 and CrebA, for RNA interference (RNAi)-mediated knockdown. Silencing either factor led to significant reductions in venom gene expression, with 13 venom genes downregulated in ds*PpATF6* treatment and 17 in ds*PpCrebA* treatment (Fig. 4e,f; supplementary Table 9,10). In both knockdown groups, the number of downregulated venom genes was greater than expected by chance (*P* < 0.0001, permutation test), whereas no such pattern was observed in the negative control (Fig. 4g-h; *P* = 0.79, permutation test). These results indicate that the contribution of these factors extends beyond a general secretory role and includes regulatory effects on venom gene expression. Taken together, our results support the idea that venom systems in parasitoid wasps and snakes share elements of regulatory logic, suggesting the parallel evolution of regulators underling venom regulation.

### Co-option of an ancestral homeobox gene as a key regulator of venom expression

Unexpectedly, we found that the Ladybird homeobox (*Lbx*) gene occupies a central position within the iGRN of venom genes and is predicted to regulate 35 venom genes (Fig. 5a; 7.73-fold enrichment, FDR adjusted *P* = 1.05e-19, one-sided hypergeometric test). *Lbx* is an ancient and highly conserved homeobox gene, traceable at least to the bilaterian ancestor. It is traditionally known for its conserved embryonic expression and roles in muscle development in diverse animal lineages, including mammals and *Drosophila*^9,33,34^. However, both bulk and single-nucleus transcriptome data reveal that *Lbx* is highly and specifically expressed in the venom gland of *P. puparum* (Tau = 0.98), with transcripts detected in 97% of venom gland cells (Fig. 5b,c; Extended Data Fig. 3a,b). Consistent with this expression pattern, the chromatin accessibility and histone modification signals showed venom gland-specific activation around the TSS of *Lbx* (Extended Data Fig. 3b). These results point to a functional co-option of *Lbx*, whereby a deeply conserved developmental regulator has acquired a derived role in venom gene regulation in parasitoid wasps. To test this hypothesis, we performed RNAi-mediated knockdown of *Lbx* followed by RNA-seq.

**Fig. 5.**
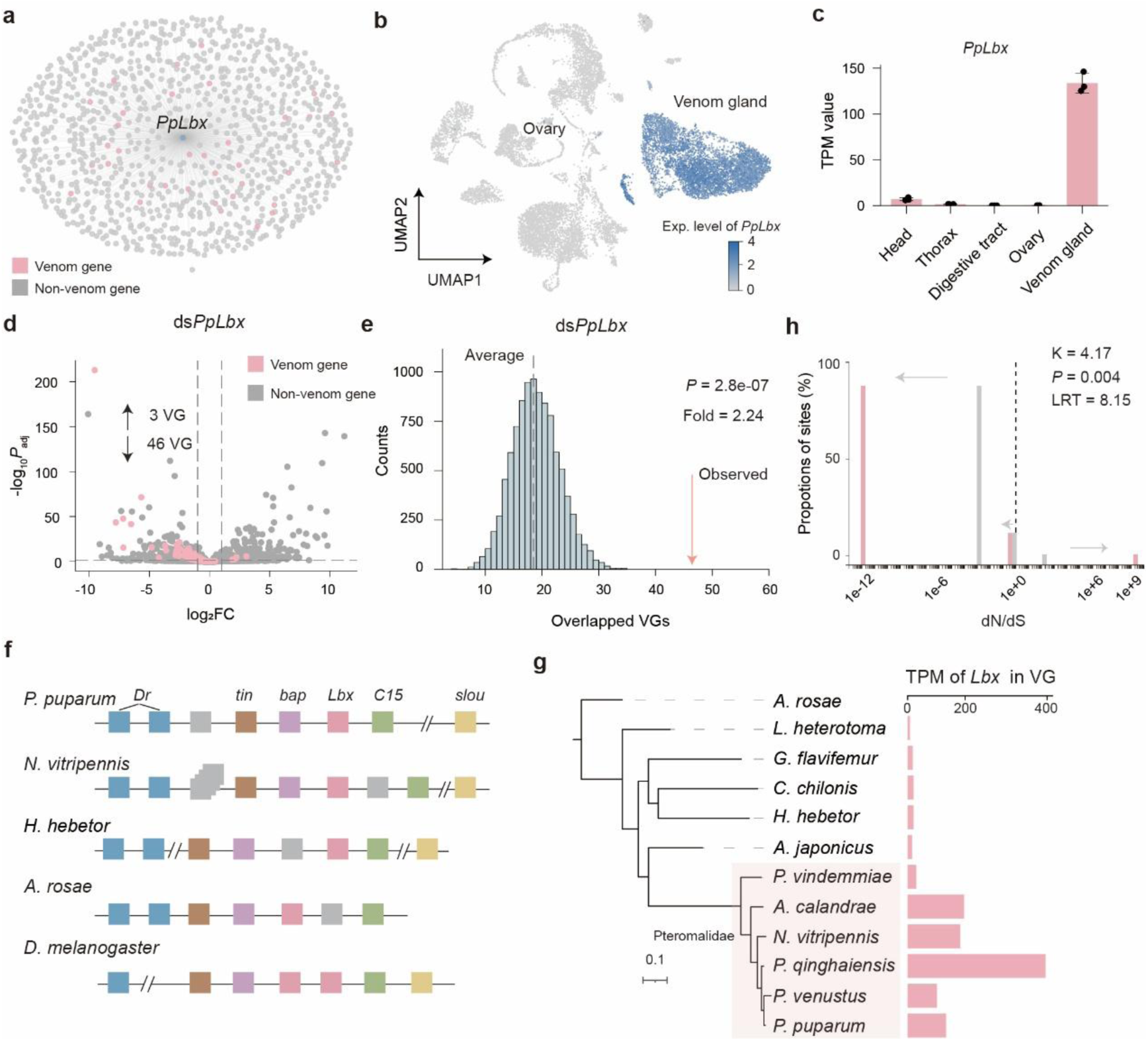
Co-option of *Lbx* into the venom gene regulatory programme in *P. puparum*. (a) *Lbx*-centred subnetwork derived from the iGRN, showing *PpLbx* and its predicted target genes, with venom genes highlighted in pink and non-venom genes in grey. (b, c) Expression profiles of *PpLbx* at both single-nucleus (b) and bulk-level (c) transctiptomes, indicating its specific and highly expression in the venom gland. (d, e) Knockdown of *PpLbx* reveals its key regulatory role in venom gene expression. *PpLbx* knockdown led to downregulation of 46 of 90 venom genes (d), a number significantly greater than expected by random sampling (e; n = 10,000). (f) Synteny analysis of the NK homeobox cluster across representative hymenopteran species and *D. melanogaster*, showing conserved genomic organization and the position of *Lbx*. (g) Phylogenetic tree of *Lbx* orthologues across Hymenoptera with venom-gland expression levels shown for each species, highlighting lineage-restricted venom-gland high expression. (h) Selection analysis of *Lbx* evolution across Hymenoptera, showing signatures of intensified natural selection in the *P. puparum*-*A. calanmdrae* clade.

Relative to dse*GFP* controls, *Lbx* knockdown did not produce obvious defects in venom gland morphology. By contrast, it caused extensive reprogramming of the venom transcriptome, with more than half of venom genes (46/90, 51%) significantly downregulated (Fig. 5d,e; Extended Data Fig. 3d-f; Supplementary Table 11; FDR adjusted *P* < 0.05, log2FC < -1). RT-qPCR independently confirmed reduced expression of these venom genes after *Lbx* knockdown, including *PpLIPK*, *PpOrcokinin*, *PpKAT3*, *PpSer6*, *PpMrjp* and *PpADA* (Extended Data Fig. 3g), establishing *Lbx* as a critical regulator of venom gene expression.

*Lbx* is a member of the NK homeobox cluster. Unlike in *Drosophila*, where *Lbx* duplicated into two paralogs, *Lbe* and *Lbl*, with distinct temporal expression patterns in early and late embryos^9^, *Lbx* has remains a single-copy gene in Hymenoptera and retains a conserved genomic arrangement within the NK clusters (Fig. 5f). Consistent with this, the *Lbx* gene tree closely mirrors the species phylogeny (Fig. 5g). To determine whether high *Lbx* expression in venom glands is conserved across parasitoid wasps, and to infer its evolutionary origin, we quantified its expression across 11 hymenopteran species using the venom gland transcriptomes. We found that elevated expression of *Lbx* in venom glands appeared specifically in a clade of pteromalid wasps (*P. puparum*-*A. calanmdrae*), which can be traced back to approximately 21 million years ago (Fig.5g). This pattern suggests that the co-option of *Lbx* into venom regulation represents a lineage-specific innovation rather than a broadly conserved feature in the evolution of parasitoid wasps. Next, we asked whether this regulatory shift was accompanied by evolutionary constraint, as venom is a key adaptive trait for parasitoid reproduction. In the clade exhibiting high venom-gland expression of *Lbx*, tests of selection detected significant intensification relative to the background (Fig. 5h; K = 4.17, *P* = 0.004, likelihood ratio test), indicating that selective pressure on *Lbx* was strengthened in this lineage. Thus, the venom-associated role of *Lbx* appears to have been selectively maintained rather than representing a transient expression change.

To further uncover the molecular basis of this co-option, we integrated ATAC-seq and histone-modification CUT&Tag data for *PpLbx* (the *Lbx* gene of *P. puparum*) and identified one active promoter and six putative active enhancers. Notably, one intronic enhancer displayed a pattern of sequence conservation that closely matched the phylogenetic origin of elevated venom-gland expression (Fig.6a,b). Dual-reporter assays confirmed that this enhancer drove strong reporter activity (Fig. 6c; *P* = 4.2e-05, one-sided *t*-test). In contrast, the orthologous sequence from *P. vindemmiae* (the closest outgroup lacking venom-gland highly expression pattern), showed no detectable activation relative to the control (Fig. 6c; *P* = 0.99, one-sided *t*-test). These results suggest that gain of a novel *cis*-regulatory element may enabled high venom-gland expression of *Lbx*, providing a mechanistic basis for the rewiring of a conserved developmental gene into an adaptive secretory network.

**Fig. 6.**
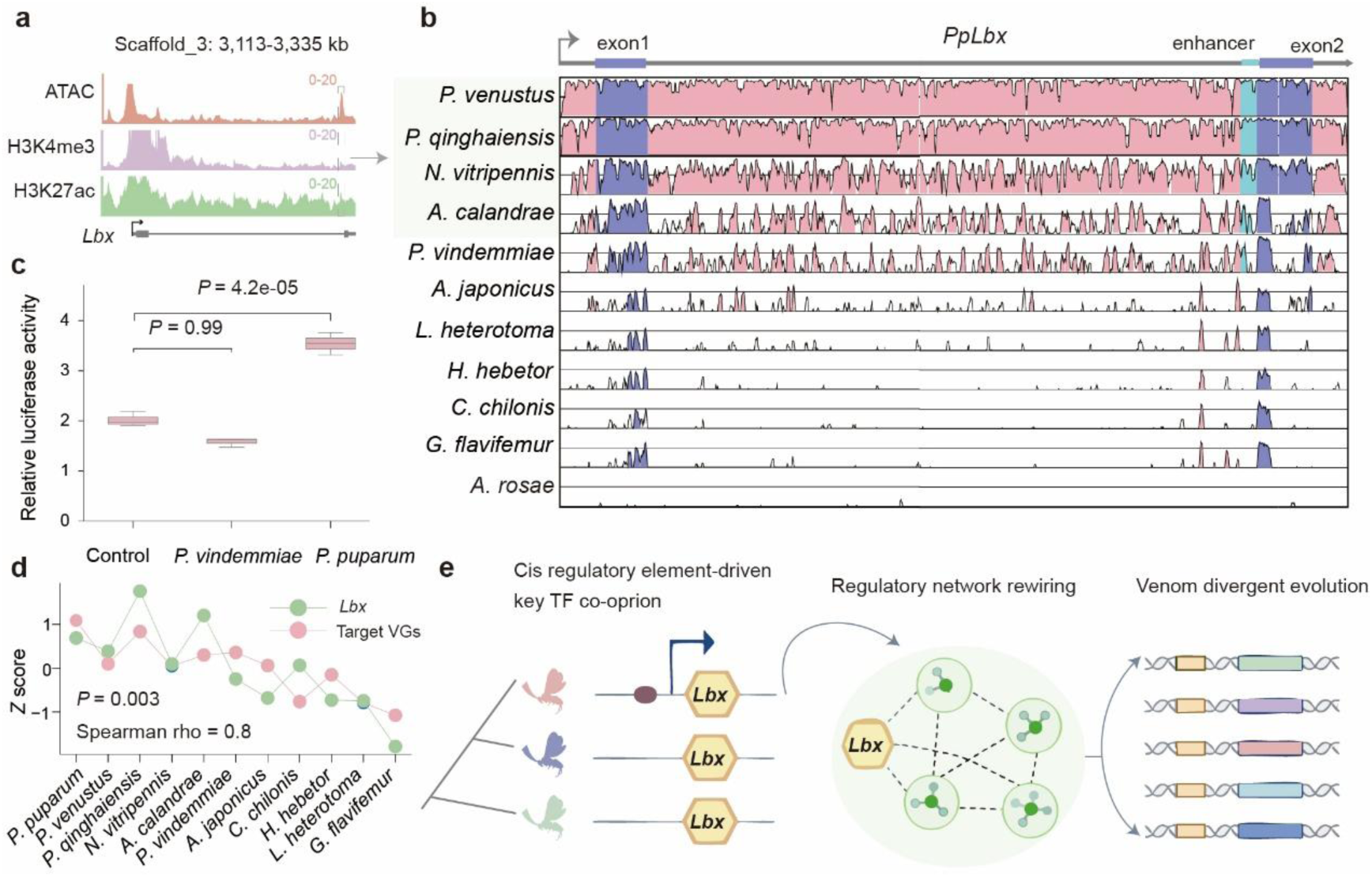
A putative intronic enhancer is associated with *Lbx* co-option in parasitoid wasps. (a) Chromatin accessibility and histone-modification profiles at the *PpLbx* locus, highlighting a putative intronic enhancer. (b) VISTA plot of sequence conservation across representative hymenopterans at the *Lbx* locus, showing that conservation of the putative enhancer broadly parallels the evolutionary pattern of *Lbx* expression. (c) Dual-luciferase reporter assays of the *P. puparum* intronic enhancer and the orthologous sequence from the closest outgroup, *P. vindemmiae*. (d) Across-species comparison of venom-gland expression (*Z* scores) of *Lbx* and its target venom genes in *P. puparum* and the corresponding orthologues in other parasitoid wasps, with Spearman’s correlation indicated. (e) Proposed model for venom evolution in parasitoid wasps, in which cis- regulatory gains co-opt key transcription factors (e.g., *Lbx*) and rewire the venom-gland gene-regulatory network, thereby driving divergent venom gene evolution.

Finally, we asked whether co-option of *Lbx* had broader consequences for venom evolution. Across parasitoid wasp lineages, the expression pattern of *Lbx* and its venom target orthologues was significantly correlated (Fig.6d; Spearman’s r = 0.9, *P* = 0.003). This indicates coordinated evolutionary change across the network. This pattern supports a model in which recruitment of *Lbx* did not simply alter a single gene, but facilitated the diversification of downstream venom repertoires through GRN-level innovation (Fig.6e).

## Discussion

Understanding how gene regulatory networks are reconfigured to generate evolutionary novelty remains a central challenge in biology^2,3,35^. Venom systems provide a powerful framework for addressing this problem because they have evolved repeatedly across animals and, in each lineage, require regulatory programmes capable of producing lineage-specific venom repertoires.^15,27,28^ Our study suggests that the origin of parasitoid venom regulation did not require the invention of an entirely new regulatory programme. Instead, the venom gland of *P. puparum* appears to have reused a conserved secretory regulatory pathway involving ER stress and UPR associated factors. This pattern appears to be conserved among parasitoid wasps and parallels regulatory features reported in snakes and other venomous lineages, including scorpions and centipedes^24,27^. Together, these findings support a general model of venom regulatory network evolution in which independently evolved venom systems repeatedly recruit an ancestral secretory regulatory framework, likely inherited from ancestral secretory cell types^27,28^.

Beyond this shared backbone, we discovered a previously unsuspected mode of innovation: the co-option of the homeobox gene *Lbx* as a central regulator of venom expression. This result was unexpected because previous developmental hypotheses suggested that venom systems derived from posterior abdominal structures, such as those of wasps or scorpions, might recruit canonical abdominal patterning regulators, including abdominal-A (*Abd-A*) gene^27^. Instead, we identified a distinct and previously unlinked homeobox factor, *Lbx*, that occupies a major regulatory position in the venom network. In its canonical context, *Lbx* is primarily associated with embryonic expression pattern, muscle precursor specification and migration, and muscle or cardiac cell identity conserved in different linages^33,36–38^. By contrast, we found that *Lbx* is specifically and strongly activated in the venom gland during the venom-production stage and is evidenced to regulate more than half of venom genes. Our epigenomic data further suggest a plausible *cis-*regulatory route for this co-option. We identified a candidate enhancer associated with *Lbx* that shows sequence patterns consistent with elevated venom-gland expression profiles across the parasitoid phylogeny. Dual-luciferase assays further confirmed its enhancer activity, whereas the orthologous sequence from a close outgroup lacking venom-gland expression showed no activity. Although direct in vivo evidence is still needed, these results support a model in which recent *cis-*regulatory changes recruited *Lbx* into the venom gland transcriptional programme. Together, these findings expand the known functional context of *Lbx* and indicate that parasitoid venom evolution involved not only reuse of a generic secretion-associated programme, but also co-option of a deeply conserved developmental regulator into a specialized secretory organ, likely through the cis-regulatory element innovation.

This framework also helps explain why parasitoid venoms appear unusually evolvable. In canonical venom systems such as snakes, venom origin and diversification often involve expansion and modification of a relatively restricted set of toxin-rich gene families, with gene duplication contributing substantially to repertoire evolution^39,40^. Parasitoid venoms, by contrast, are characterized by rapid turnover, extensive lineage-specific gains and losses, and mainly through co-option of a large catalogue of genes that were not ancestrally venom-biased^21,22,34^. A regulatory architecture centered on a co-opted homeobox factor provides a mechanistic explanation for this contrast. As homeobox transcription factors occupy high positions in developmental hierarchies and can coordinate large downstream gene sets^41,42^, their recruitment into the venom gland would create an efficient chance for recruitment of diverse genes into the venom repertoire. In this view, the conserved secretory module supplies the foundation for venom regulatory, while *Lbx* provides the regulatory leverage needed for the unusually flexible and large-scale gene recruitment that distinguishes parasitoid venoms (Fig. 6e).

More broadly, our results represent a case of homeobox gene functioning beyond canonical development, thereby expanding our understanding of homeobox gene function and evolution. Since their discovery in *Drosophila* in the 1980s, homeobox genes have been viewed as deeply conserved regulators of body plan patterning and positional identity^10,43^. Recent studies have shown that members of this ancient family can be redeployed in new contexts to generate phenotypic novelty^10^, for instance in wing evolution^5,44^, newly evolved structures like genital structures in *Drosophila*^7^, horns in beetles^45^, silk production organ in spiders^46^, and lighting organs in the firefly^47^. Most of these examples, however, remain tied to the developmental construction or modification of morphological traits. In contrast, the post-developmental expression and function of homeobox genes remain much less explored, with only a few cases functionally characterized, mainly in model organisms^10^. In *Drosophila*, for instance, adult expression of the homeobox gene (*Ubx*) in the nervous system has been linked flight behavior^48^. Against this background, our findings reveal a particularly clear example of post-developmental homeobox gene co-option in an adaptive tissue. We showed that *Lbx* has been co-opted into the adult venom system of parasitoid wasps, where it does not pattern venom gland development but instead regulates venom production. Thus, this gene appears to have been recruited from a canonical developmental framework into a physiological and ecological function, expanding the conventional view of homeobox gene co-option and providing new insights into their functional divergence during evolution.

In summary, our findings reveal that the parasitoid venom GRN is built from two nested layers: a conserved ER stress and UPR response backbone and lineage-specific rewiring through the co-option of a developmental homeobox regulator into the venom program. Moreover, our work shows that homeobox gene evolution is not limited to the redeployment of developmental regulators in new patterning or morphological contexts. Instead, a deeply conserved homeobox gene can be co-opted into a mature physiological tissue, where it regulates the production of secreted effectors. This unexpected post-developmental function of *Lbx* expands the known functional landscape of homeobox genes and provides new insight into how developmental regulators acquire physiological roles during evolution.

## Methods

### Animals and sample collection

*P. puparum* and its host, *Pieris rapae*, were reared under laboratory conditions at 25 °C and 50% relative humidity under a 16 h light/8 h dark cycle. For tissue-resolved profiling of transcriptional and epigenomic landscapes in *P. puparum*, venom glands, ovaries, digestive tracts, thoraxes, and heads were dissected from 2-day-old adult females of the laboratory strain Ppup_zju and subjected to downstream multi-omic sequencing. Nuclei isolated from each tissue were used as input for ATAC and CUT&Tag sequencing library construction and prepared as previously describe^49,50^. Briefly, tissue samples were collected and homogenized in homogenization buffer (containing 0.1% Triton X-100 and 0.1 mM DTT), filtered through a 35-μm cell strainer, washed, and centrifuged at 500g for 5 min at 4 °C to prepare for subsequent library construction.

### ATAC-seq

Paired-end ATAC-seq libraries were generated using the Hyperactive ATAC-Seq Library Prep Kit for Illumina (TD711, Vazyme) according to the manufacturer’s instructions. Briefly, ∼50,000 nuclei were isolated from each tissue sample as described above and resuspended in 50 μl transposition reaction buffer containing 0.1% Tween-20, 0.01% digitonin, 1× TTBL and 0.1% TTE Mix V50. Transposition was performed at 37 °C for 30 min with gentle mixing. The reaction was terminated by adding 5 μl STOP buffer, and released DNA fragments were purified using ATAC DNA Extract Beads. Libraries were amplified by PCR with indexed N5 and N7 primers and sequenced on the Illumina NovaSeq platform. Three biological replicates were generated for each tissue.

ATAC-seq data were processing using the ENCODE ATAC-seq pipeline v2.3.3 (https://github.com/ENCODE-DCC/atac-seq-pipeline). Briefly, raw reads were adapter and quality trimmed using cutadapt^51^ v1.9.1, aligned to the *P. puparum* reference genome using Bowtie2^52^ v2.2.9 with --very-sensitive -X 1000 --dovetail paramaters. Reads mapped to the mitochondrial genome were removed, and only alignments with mapping quality scores of at least 20 were retained. PCR duplicates were marked and removed using Picard v2.10.6 (https://github.com/broadinstitute/picard). Peaks were called with MACS2^53^ v2.1.1 using the parameters --nomodel --shift -100 --extsize 200. Reproducible peaks across biological replicates were identified using IDR^54^ v2.0.4.2, and the conservative IDR peak set was retained for downstream analyses. Quality control metrics, including fragment size distribution, TSS enrichment, FRiP and library complexity, were assessed within the ENCODE pipeline and cisDynet^55^ v1.0.0. Transcription factor footprinting analysis was conducted using the TOBIAS^56^ Snakemake workflow (https://github.com/loosolab/TOBIAS_snakemake) with default parameters, including ATACorrect for Tn5-bias correction, ScoreBigwig for footprint scoring and BINDetect for motif occupancy analysis.

### CUT&Tag

CUT&Tag libraries for H3K4me3 and H3K27ac profiling in the five tissues were generated using the Hyperactive Universal CUT&Tag Assay Kit for Illumina Pro (TD904, Vazyme) according to the manufacturer’s instructions. Approximately 50,000 nuclei were prepared for each sample and immobilized on conA beads at room temperature for 10 min. Bead-bound nuclei were incubated with primary antibodies against H3K4me3 (Cat. No. 39160, Active Motif) and H3K27ac (Cat. No. 39134, Active Motif) overnight at 4 °C. Samples were then incubated with a secondary antibody (Cat. No. Ab207-01, Vazyme) for 1 h at room temperature, washed twice, and incubated with pA/G-Tn5 Pro at room temperature for 1 h. Tagmentation was performed in TTBL buffer at 37 °C for 60 min. DNA was extracted using DNA Extract Beads Pro, amplified by PCR with indexed N5 and N7 primers, and purified using VAHTS DNA Clean Beads. Final libraries were quantified using a Qubit3 and assessed on an Agilent Bioanalyzer, and then sequenced using the Illumina NovaSeq platform. Three biologically independent replicates were generated for each tissue.

CUT&Tag data were processed using nf-core/cutandrun pipeline v3.2.2^57^. Raw reads were filtered using Trim Galore v0.6.6 (https://github.com/FelixKrueger/TrimGalore), aligned to the *P. puparum* reference genome with Bowtie2^52^ v2.4.4, and filtered to remove low-quality (MAPQ <20), unmapped, mitochondrial and duplicate reads. Peaks were called using MACS2^53^ v2.2.7.1 with patermters -q 0.01 -f BAMPE -keep-dup all. Consensus peaks reproducibly detected across biological replicates were retained based on IDR^54^ analysis. Peak annotation was performed using ChIPseeker^58^ v1.32.1.

### RNA-seq

Total RNA was isolated from each tissue using RNAiso Plus (Cat. No. 9109, Takara) and used for library preparation with the VAHTS Universal V10 RNA-seq Library Prep Kit for Illumina (NR606-01, Vazyme). Libraries were sequenced on the Illumina NovaSeq platform. Three biologically independent replicates were generated for each tissue.

Raw RNA-seq reads were filtered using Trim Galore v0.6.6 (https://github.com/FelixKrueger/TrimGalore). Clean reads were aligned to the *P. puparum* reference genome using HISAT2^59^ v2.2.1. The expression level at both transcript and gene level were quantified and summarized to TPM using salmon^60^ v1.10.3. Differential expression analysis was performed on raw gene-count matrices using PyDESeq2^61^ v 0.5.3. Genes with an FDR adjusted *P* value <0.05 and |log2 fold change| >1 were considered differentially expressed. To identify tissue-specific genes, we calculated the tissue-specificity index Tau using tspex^62^ v0.6.3 for each gene across the five tissues. Genes with Tau >0.8 were classified as tissue-specific, and the tissue showing the highest TPM value was assigned as the representative tissue of specificity.

### Identification and evolutionary analysis of putative active cis-regulatory elements

Active cis-regulatory elements were inferred by integrating chromatin accessibility with active histone marks. Putative active promoters (pAPs) were defined as accessible regions overlapping both H3K4me3 and H3K27ac peaks, and putative active enhancers (pAEs) were defined as accessible regions overlapping H3K27ac peaks but lacking detectable H3K4me3 enrichment signals. To investigate the evolutionary conservation of pAPs and pAEs, and its potential roles in venom gene evolution. We selected 14 additional represent parasitoid wasp species, which have high quality genome data and available transcriptome data of venom gland (Supplementary Table 7). We constructed a 15-way whole-genome multiple alignment using Progressive Cactus^63^ v 2.9.9, using the species tree adopted from previous published parasitoid phylogeny as a guild tree^20^. For each pAP or pAE in *P. puparum* genome, the corresponding aligned region was extracted from the whole-genome alignment and evaluated for alignment coverage. Elements with alignment coverage ≥80% were classified as conserved. As a control, the exonic regions of the related venom genes were analysed using the same procedure. To assess the relationship of the evolution of pAPs and pAEs and venom gene expression profiles, we inferred orthologous gene relationships using the TOGA v2.0 pipeline^64^. Gene expression levels were quantified from RNA-seq data following the previously described workflow. We retained only single-copy orthologous genes and constructed a TPM matrix for the corresponding orthologous groups. TPM values were log2-transformed followed by quantile normalization and *Z*-score scaling within each species. The resulting values represent the relative expression intensity of each gene in the venom gland within a given species.

### Transcription factors identification

All protein sequences in *P. puparum* genome were screened using InterProScan^65^ v6.0 and searched against AnimalTFDB^66^ v4.0 with BLASTP 2.17.0. Proteins were considered candidate TFs if they contained a DNA-binding domain according to DBD database^67^ and showed significant similarity to a known TF. We then classified each TF according to the AnimalTFDB classification scheme and manually curated the results. To infer the potential binding motifs of these TFs, we applied the JASPAR profile inference tool^68^ and assigned putative known motifs to each TF, yielding 813 motifs corresponding to 184 TFs.

### Construction of gene regulatory network

We reconstructed the GRN using an integrative framework that combines five individual networks, capturing distinct physical and functional features^69–71^. The two physical networks comprised a motif-based regulatory network (PWM) and a footprinting-based regulatory network. For the PWM network, we scanned promoter regions (-1500 to +500 bp relative to TSS) of each gene using FIMO^72^ v5.5.9 with inferred motifs of *P. puparum* TFs, and used motif-matching scores as interaction weights. This analysis yielded 181 TFs connected by 2,659,712 putative TF-target interactions. For the footprinting network, we analyzed our ATAC-seq data as described above to infer TF occupancy footprints. A TF-target interaction was defined when a footprint-associated binding event was detected in the regulatory region of the target gene, and the corresponding binding score was used as the interaction weight. This network contained 184 TFs and 853,595 interactions. The three functional networks included two co-expression networks (CoE and GENIE3) and one co-modification network (CoC). To construct the co-expression-based networks, we combined newly generated RNA-seq data with publicly available transcriptomic datasets, resulting in a compendium of 85 transcriptomes spanning multiple developmental stages and tissues. For the CoE network, Spearman correlation coefficients between each TF and gene were calculated, retaining the top 1,500 associations per TF and using correlation as interaction weight. This network comprised 754 TFs and 1,131,000 interactions. For the second co-expression network, we applied GENIE3^73^, which uses tree-based ensemble learning to capture potentially non-linear regulatory relationships, yielding 754 TFs and 857,473 interactions. The co-modification network (CoC) was constructed by quantifying the correlation of chromatin modification states between TFs and potential target genes, resulting in 711 TFs and 1,066,500 interactions.

Next, the five individual networks were integrated into a comprehensive GRN using a supervised machine-learning framework. Interaction weights from each input network were first rescaled to a range of 0-1 using previous described^71^. The positive training set was derived from the REDfly^74^ regulatory network (experimentally validated interactions), retaining only TF-target pairs with single-copy orthologues in *P. puparum*, yielding 340 positive interactions. The remaining corresponding TF-target pairs involving TFs represented in the positive set were treated as putative negatives, from which negative samples were randomly drawn at a 3:1 ratio relative to positives. The dataset was randomly split into training and test sets at a ratio of 8:2. We then evaluated multiple binary classification models, including logistic regression, random forest, support vector classification, and gradient boosting, implemented with the scikit-learn package in Python. Logistic regression showed the best overall performance and was therefore selected for final prediction (Supplementary Table 12). The resulting integrated GRN comprised 731 TFs and 456,902 high-confidence regulatory interactions. Since a gold-standard regulatory dataset is not yet available for *P. puparum*, we additionally applied an unsupervised integration strategy as an independent validation step. For each TF-target pair, weights across the individual input networks were summed, and interactions within the top 10% of summed scores were retained to construct an unsupervised network. The supervised and unsupervised approaches showed 78% concordance, and interactions consistently supported by both methods were retained as the final integrated GRN, comprising 718 TFs and 361,466 interactions.

### Evaluation and analysis of regulatory network

To assess whether the iGRN captured complementary information and outperformed individual networks, we first quantified the overlap of interactions between the iGRN and each input network. Network contribution and integration were measured as the proportion of shared interactions relative to each input network and to the iGRN, respectively. We next evaluated the enrichment of interactions from the iGRN and each independent input network in the evaluation test dataset. Fold enrichment was calculated as the ratio of the observed number of overlapping interactions to the expected number of overlaps under a random interaction model.

Next, we used orthologous ChIP-seq datasets from *Drosophila* for independent validation^75^. Only TF-target pairs conserved as orthologues in *P. puparum* were considered. The observed overlap was compared to a null distribution generated by 10,000 randomizing network edges while preserving network topology, controlling for structural biases. Additionally, we evaluated the iGRN based on the functional co-occurrence of targets sharing common regulators, on the premise that co-regulated genes are more likely to exhibit similar expression profiles and biological functions. We first identified gene pairs with a Jaccard similarity score greater than 0.5 as co-regulated pairs. Pairwise Pearson correlation coefficients were then calculated for the expression profiles of these co-regulated gene pairs, and GO similarity was assessed using the GO3^76^ package. The observed distributions were compared with those derived from 10,000 randomized networks, confirming that the iGRN captured functional regulatory coherence beyond random expectation.

### Evolutionary analysis of *Lbx*

The structure of *Lbx* is highly conserved, and typically contains a long intron (e.g. ∼17 Kb in *P. puparum*) in parasitoid wasp. However, automated genome annotation pipelines likely fragmented this gene model. To accurately define *Lbx* gene structures, we first identified candidate loci in each genome assembly by aligning the *Drosophila Lbx* to the genomes using tBLASTN. Putative loci were then extended by 5 kb upstream and downstream and subjected to gene prediction using FGENESH+^77^. The resulting gene models were validated using RNA-seq data from each species (Supplementary Table 7) to ensure concordance with expressed transcripts.

The single-nucleus expression profiles of *PpLbx* in the venom gland and ovary were obtained from our previous dataset and visualized using Scanpy^50^. The gene synteny analysis of the NK cluster homeobox genes were performed based on the orthologues relationship inferred by OrthoFinder^78^ v3.0.1b1. We then reconstructed the phylogeny of *Lbx* using IQ-TREE^79^ v3.0.1 with 1,000 bootstrap replicates, based on the nucleotide alignment generated by MAFFT^80^ v7.525 and the best-fit substitution model selected by ModelFinder^81^. The resulting phylogenetic tree was annotated with expression data, which were quantified using salmon^60^ v1.10.3 (as described above) and mapped onto the tree to reveal evolutionary and regulatory patterns. To test whether selective pressure on *Lbx* was altered in the *P. puparum*-*A. calanmdrae* clade, we applied RELAX^82^ implement in the Hyphy^83^, which estimates a selection-intensity parameter, K. A significant result of k>1 indicates that selection strength has been intensified along the test branches, and a significant result of k<1 indicates that selection strength has been relaxed along the test branches. The significance of both models was evaluated using a likelihood ratio test.

### RNAi and RNA-seq of venom glands

To characterize the roles of *PpAtf6*, *PpCrebA* and *PpLbx* in regulating venom gene expression, we performed RNAi-mediated knockdown followed by RNA-seq of venom glands. Briefly, gene-specific primers (Supplementary Table 13) were designed to amplify a 300-500 bp fragment of each target gene, and the resulting amplicons were verified by Sanger sequencing. Purified DNA fragments of each target genes together with the control gene (*eGFP*), were used as templates for dsRNA synthesis using the T7 RNAi Transcription Kit (Cat. No. TR102, Vazyme) according to the manufacturer’s instructions. The dsRNAs were purified using RNA Clean Beads, eluted in nuclease-free water, and quantified using a NanoDrop 2000 (Thermo Fisher Scientific). Their integrity and size were further verified by electrophoresis on a 1% agarose gel. dsRNA injection in *P. puparum* was performed as described previously^84^. Briefly, yellow female pupae derived from the offspring of a single adult female were collected and randomly assigned to three treatment groups and one dse*GFP* control group. Approximately 200 ng dsRNA was injected into the abdomen of each yellow pupa using a nano-injector.

Venom glands were dissected from 2-day-old adult females from each group, and RNA-seq libraries were constructed and then sequenced on the MGI platform. Each group included three to six biological replicates. Transcriptome data processing, expression quantification, and differential expression analysis were performed as described above. Principal component analyses of both the global transcriptome and the venom-gene expression matrix were performed using PCAtools (https://github.com/kevinblighe/PCAtools). To determine whether venom genes were disproportionately downregulated after knockdown, we compared the observed number of downregulated venom genes with the number expected by chance and assessed significance using Fisher’s exact test.

### Quantitative real-time PCR analysis

Total RNA was isolated using a RNAiso Plus (Cat. No. 9109, Takara) and reverse transcribed into cDNA using HiScript III RT SuperMix for qPCR (Cat. No. R231, Vazyme) following the manufacturer’ s instructions. qPCR was performed with gene-specific primers (Supplementary Table 13) on a LightCycler 480 II System (Roche Diagnostics) using ChamQ SYBR qPCR Master Mix (#Q321-02, Vazyme). Relative expression was calculated using the 2-^ΔΔCt^ method^85^, and statistical analyses were performed in Python with SciPy.

### Dual Luciferase reporter assay

The putative enhancer of *Lbx* in *P. puparum*, and its orthologues sequences from the outgroup *P. vindemmiae* were chemical synthesized, cloned into the pGL4.23[luc2/minP] vector (Promega), and verified by Sanger sequencing. Details of the inserted fragments are provided in Supplementary Table 14. Reporter plasmids were transiently transfected into HEK293T cells together with the Renilla luciferase control plasmid pGL4.74[hRluc/TK] for normalization using Lipo8000™ Transfection Reagent (Beyotime) according to the manufacturer’s instructions. After 36h, cells were lysed, and firefly and Renilla luciferase activities were measured using the Dual-Luciferase Reporter Assay System (Promega). Relative activity was calculated as the ratio of firefly luciferase to Renilla luciferase activity.

### Enrichment analysis

GO annotations for the protein sequences were generated using eggNOG-mapper online serves^86^ with default parameters. Enrichment analyses were performed using the clusterProfiler^87^ package. Terms with an adjusted *P* value < 0.05 were considered significantly enriched.

## Data availability

The raw sequencing data generated in this study, including RNA-seq, ATAC-seq, and CUT&Tag sequencing reads has been deposited in the Genome Sequence Archive at the National Genomics Data Center of the China National Center for Bioinformation under the BioProject accession number PRJCA064322.

## Acknowledgements

This work was supported by the Program of National Natural Science Foundation of China (NSFC) (grant No. 32472631 and 32202376 to X.Y.; Grant No. 32302428 to Y.Y.), Key Program of NSFC (Grant No. 32330085 to G.Y.), the Scientific Research Startup Fund Project of Zhejiang Agriculture and Forestry University (Grant No. 2023LFR119 to X.Y.).

## Contributions

X.H.Y., G.Y.Y. and Y.Y. conceived, designed and supervised the study. Y.Y., S.S.W., C.L. and S.X.L. collected the samples and prepared the sequencing libraries. Y.Y. and S.S.W. analysed the data. Y.Y., S.S.W., S.X., Z.C.C. and Y.Y.C. performed the experiments. Y.Y. and X.H.Y. wrote the manuscript, with critical input from Q.F. and G.Y.Y. All authors reviewed and approved the final manuscript.

## Ethics declarations

### Competing interests

The authors declare no competing interests.

## Figures

**Extended Data Fig. 1.**
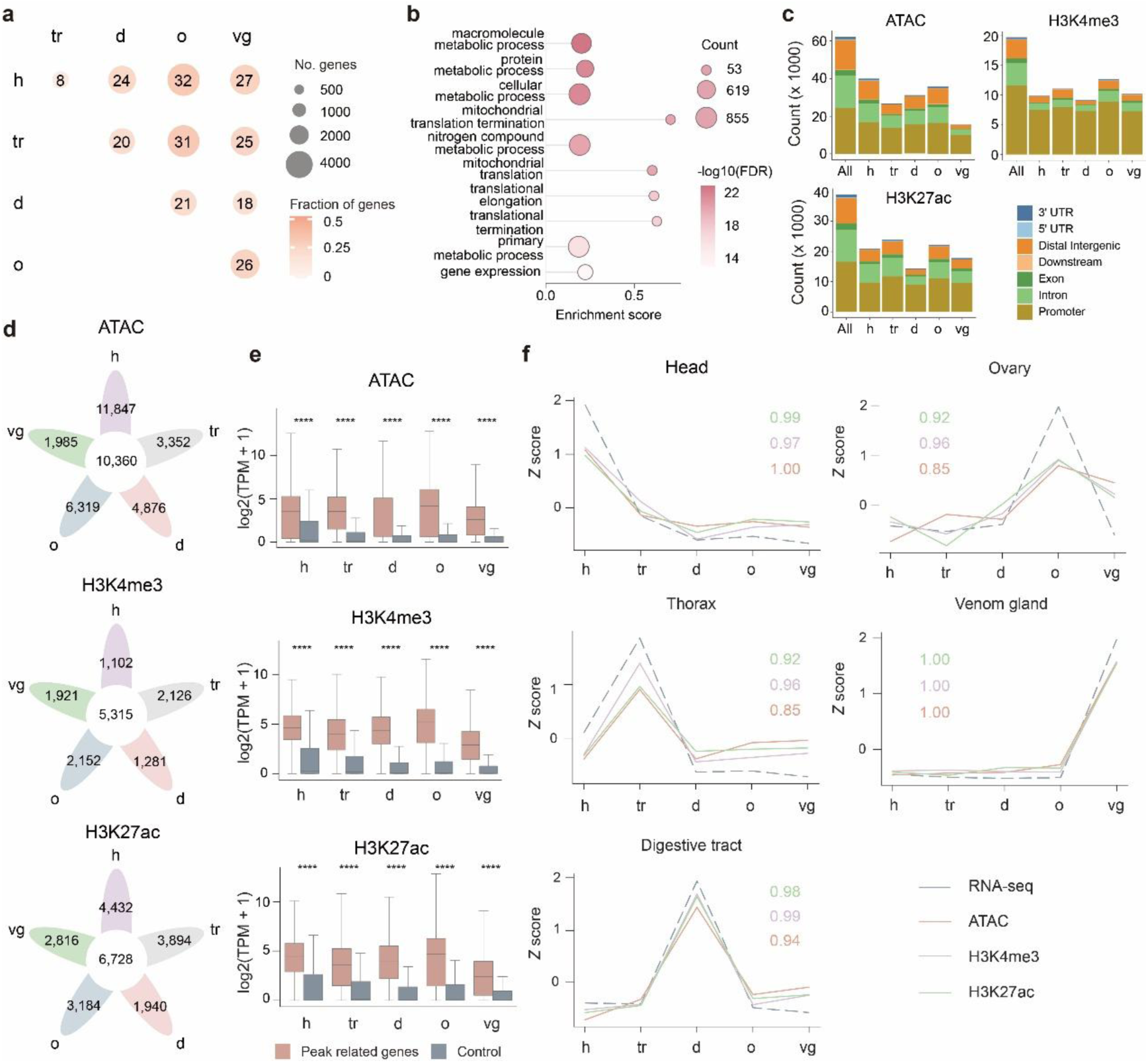
Tissue-specific transcription is coupled to active chromatin states in *P. puparum.* (a) Pairwise differential expression analysis across five tissues: head (h), thorax (tr), digestive tract (d), ovary (o) and venom gland (vg). (b) Gene Ontology enrichment analysis of constitutively expressed genes. (c) Genomic distribution of ATAC-seq, H3K4me3 and H3K27ac peaks across individual tissue peak sets and merged sets. (d) Overlap of ATAC-seq, H3K4me3 and H3K27ac peaks among tissues. Numbers indicate peak counts in tissue-specific or shared categories. (e) Expression levels of genes associated with ATAC-seq, H3K4me3 or H3K27ac peaks compared with control genes lacking the corresponding peaks. **** indicate *P* < 0.0001, one-sided Mann-Whitney *U* test. (f) Correlation between tissue-specific gene expression and chromatin signals. For genes specifically expressed in the indicated tissue, *Z* score-normalized RNA-seq expression and ATAC-seq, H3K4me3 and H3K27ac signals are shown across tissues. Numbers indicate Pearson’s correlation coefficients between RNA-seq expression and each chromatin signal.

**Extended Data Fig. 2.**
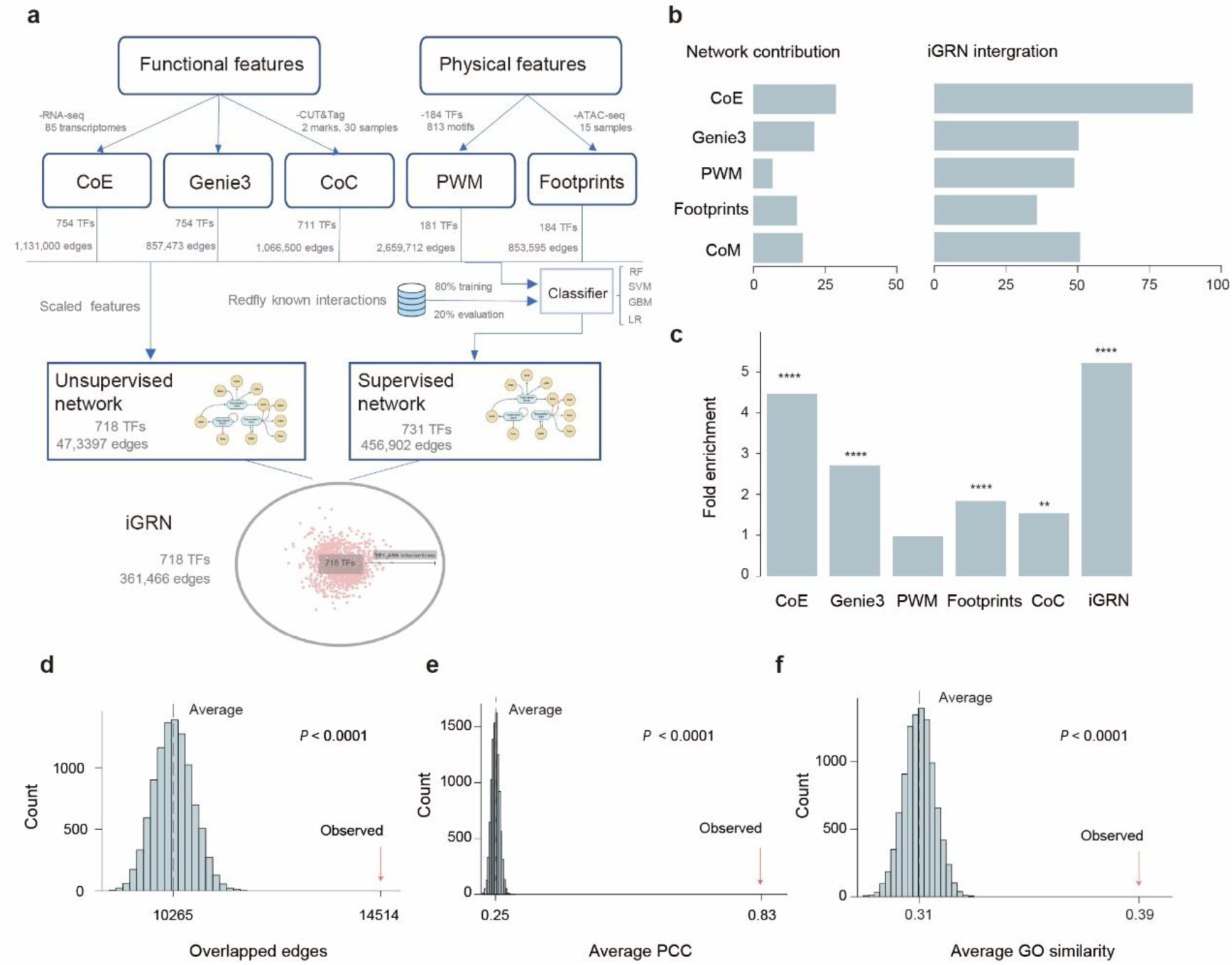
Constructing and evaluation of the iGRN. (a) Overview of the five initial input networks and the integration strategy used to construct the iGRN. Functional feature-based networks include co-expression (CoE), GENIE3 and co-chromatin activity (CoC) networks. Physical feature-based networks include motif-based PWM predictions and ATAC-footprint-based networks. (b) Complementary contributions of independent input networks to the iGRN. Bars on the left indicate the fraction of each input network retained in the iGRN, and bars on the right indicate the fraction of iGRN interactions supported by each input network. (c) Fold enrichment of the iGRN and each independent input network for interactions in the evaluation test dataset. **** *P* < 0.0001, ** *P* < 0.01, one-sided hypergeometric test. (d-f) Benchmarking of iGRN performance using independent validation datasets. The iGRN was evaluated by enrichment for interactions supported by *Drosophila* orthologue ChIP-seq datasets (d), similarity in gene expression profiles between co-regulated gene pairs (e) and similarity in Gene Ontology annotations between co-regulated gene pairs (f). Null distributions were generated by network randomization (n = 10,000) while preserving network structure. Red arrows indicate observed values, and *P* values were calculated by permutation tests.

**Extended Data Fig. 3.**
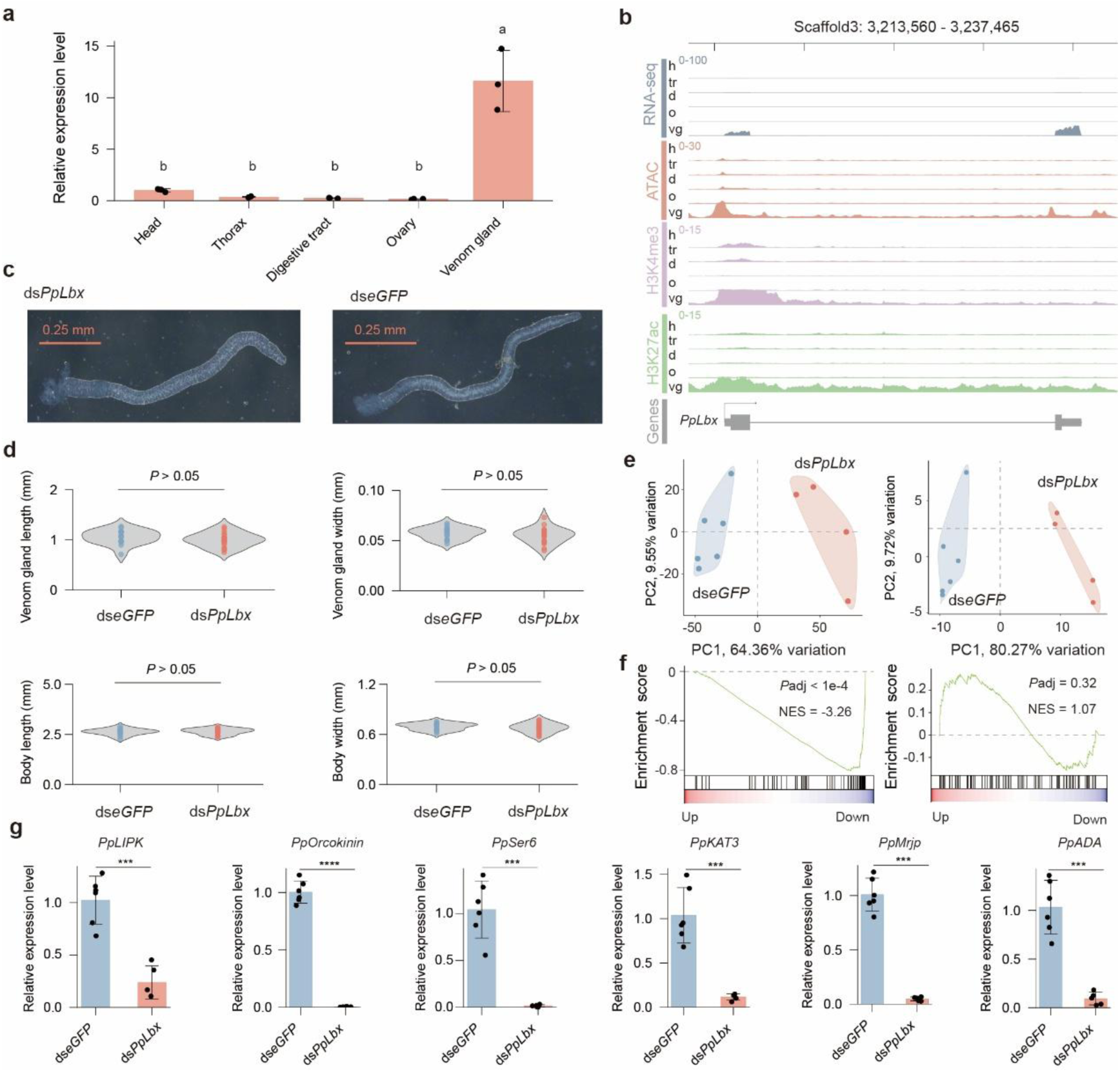
The expression profile and functional analysis of *PpLbx.* (a) Tissue expression profile of *PpLbx* measured by qRT-PCR. Error bars indicate s.d., and letters above bars denote Tukey multiple-comparison groups; different letters indicate *P* < 0.05. (b) Genome-browser tracks showing RNA-seq, ATAC-seq, H3K4me3 and H3K27ac signals at the *PpLbx* locus across five tissues. Venom gland-preferred RNA-seq and active chromatin signals are observed around the *PpLbx* transcription start site. (c, d) Morphological comparison of venom glands after *PpLbx* knockdown. Representative venom glands from ds*PpLbx* and ds*eGFP* treated females are shown in (c). Quantification of venom gland length and width, as well as body length and thorax width, is shown in (d). No significant differences were detected between ds*PpLbx* (n = 18) and ds*eGFP* (n = 15) groups. *P* values were calculated using Mann-Whitney *U* tests. (e) Principal component analysis of RNA-seq samples from ds*PpLbx* and ds*eGFP* treated venom glands based on all expressed genes (left) and venom genes (right). Samples separate clearly by treatment group in both analyses, indicating that *PpLbx* knockdown alters the venom gland transcriptome. (f) Gene set enrichment analysis of genes differentially expressed after *PpLbx* knockdown. Genes downregulated after *PpLbx* knockdown are significantly enriched among venom genes(left), but not among randomly selected control gene sets (right). NES, normalized enrichment score. (g) qRT–PCR analysis of selected venom genes downregulated after *PpLbx* knockdown. Bars show relative expression levels in ds*PpLbx* treated samples compared with ds*eGFP* controls. Error bars indicate s.d. Asterisks indicate statistical significance determined by two-tailed independent-samples t-tests; **** *P* < 0.0001; *** *P* < 0.001.

## Notes

### Competing Interest Statement

The authors have declared no competing interest.

