## Supplementary Figures for "Regulatory co-option of a homeobox gene drives parasitoid venom evolution"

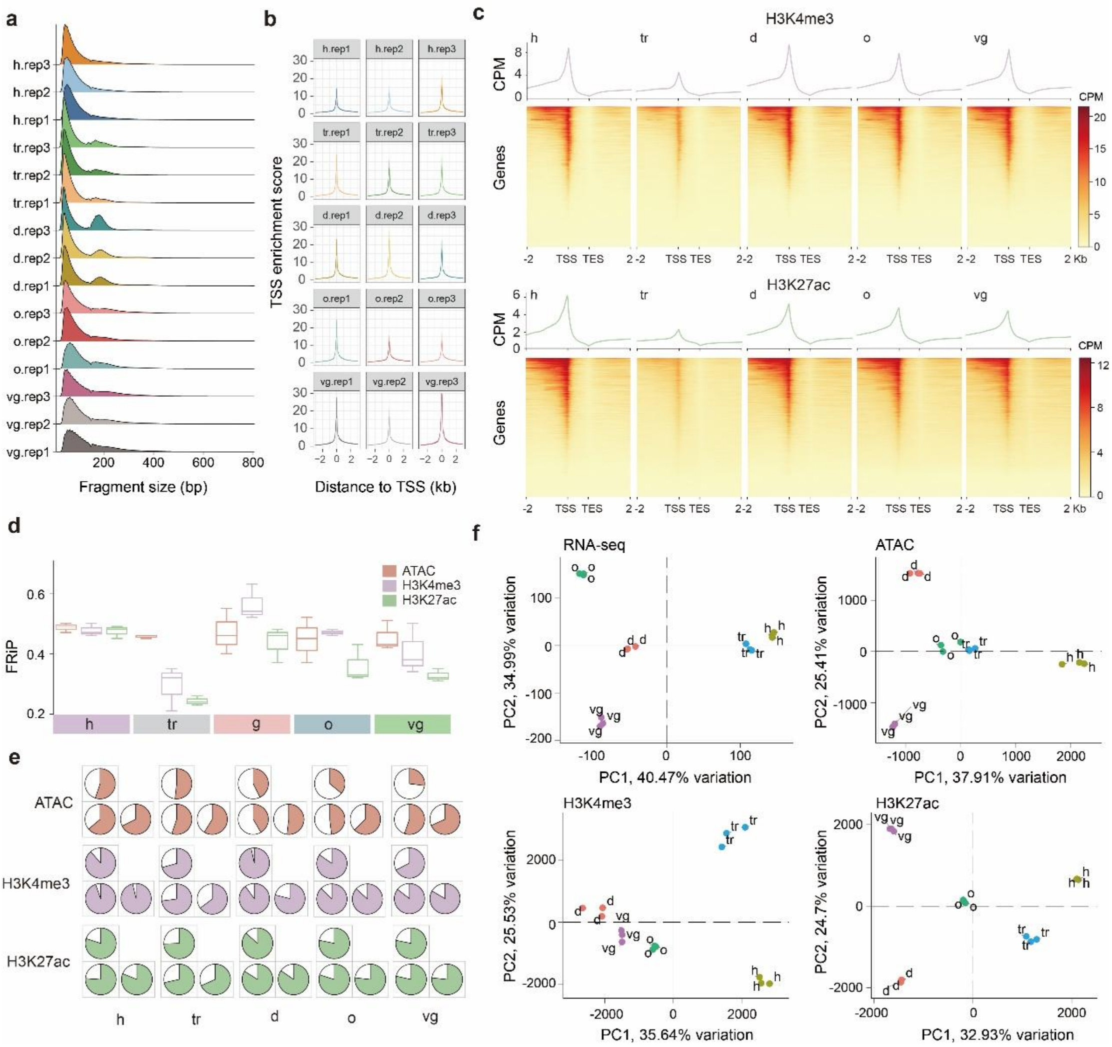


Supplementary Fig. 1 Quality control of ATAC-seq and CUT&Tag datasets. (a-b) Quality control of ATAC-seq libraries. Fragment-size distributions and transcription start site (TSS) enrichment profiles are shown for each ATAC-seq sample. (c) Heatmaps and average profiles showing H3K4me3 and H3K27ac signals across gene bodies and flanking regions in each tissue. Signals are centred from the TSS to the transcription end site (TES), with 2-kb upstream and downstream regions shown. (d) Fraction of reads in peaks (FRiP) for ATAC-seq, H3K4me3 CUT&Tag and H3K27ac CUT&Tag libraries across tissues. (e) Peak-overlap analysis between biological replicates for ATAC-seq, H3K4me3 CUT&Tag and H3K27ac CUT&Tag datasets. (f) Principal component analysis of RNA-seq, ATAC-seq, H3K4me3 CUT&Tag and H3K27ac CUT&Tag samples. Biological replicates cluster together, supporting the reproducibility of the sequencing datasets.

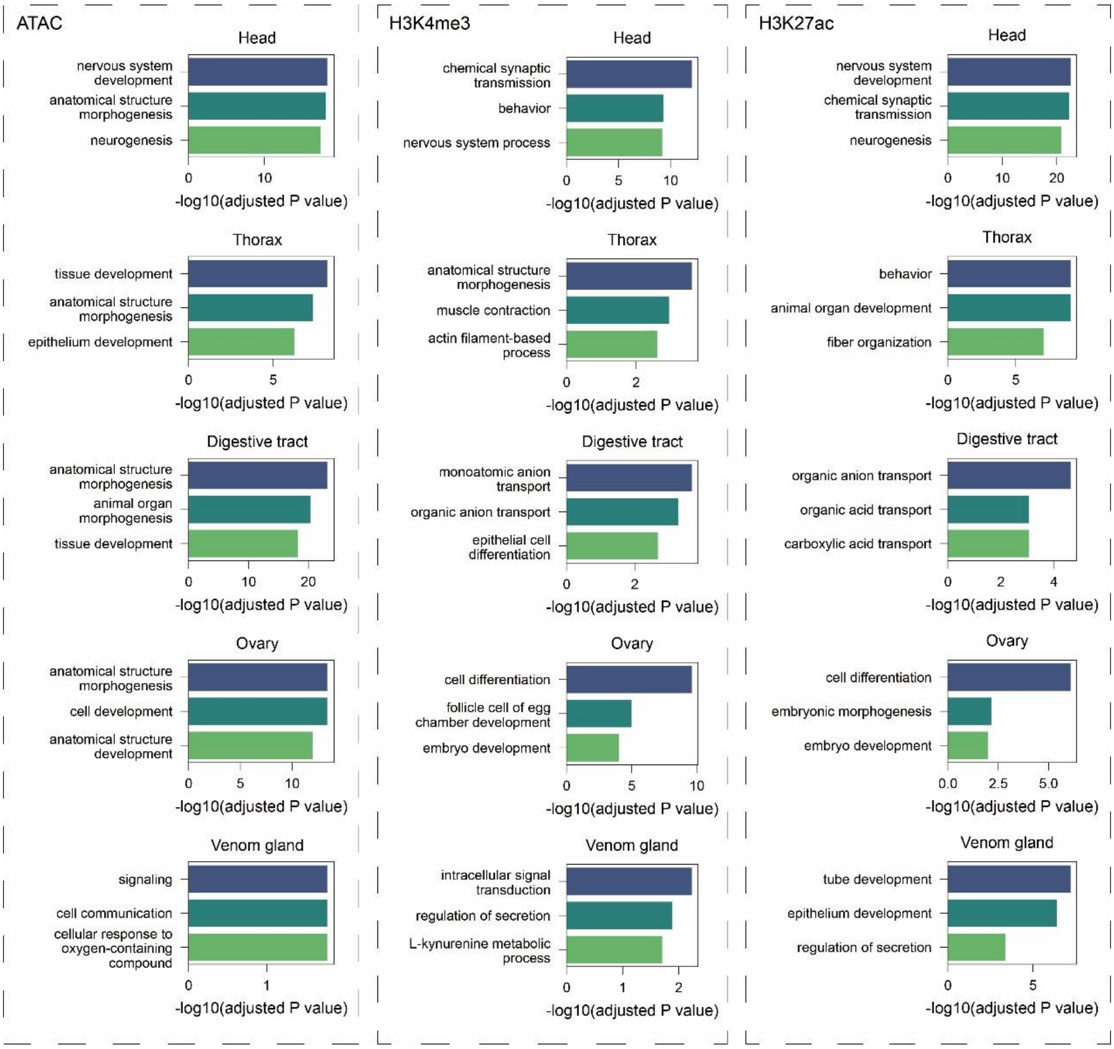


Supplementary Fig. 2 Gene Ontology enrichment of genes associated with tissue-specific chromatin peaks. Gene Ontology enrichment analysis of genes associated with tissue-specific ATAC-seq, H3K4me3 and H3K27ac peaks in the head, thorax, digestive tract, ovary and venom gland. For each tissue and chromatin feature, the top enriched GO biological process terms are shown.

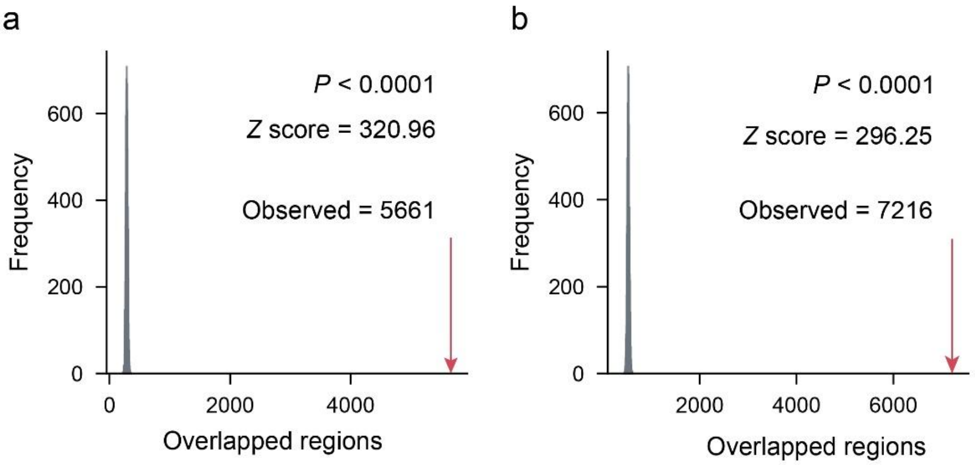


Supplementary Fig. 3 Significant co-occurrence of ATAC-seq peaks with H3K4me3 and H3K27ac peaks. a, b, Permutation analysis (n = 10,000) of the overlap between ATAC-seq peaks and H3K4me3 (a) or H3K27ac (b) CUT&Tag peaks. Grey distributions indicate the expected numbers of overlapped regions from randomized peak sets. Red arrows indicate the observed numbers of overlapped regions. *P* values and Z scores were calculated from permutation tests.

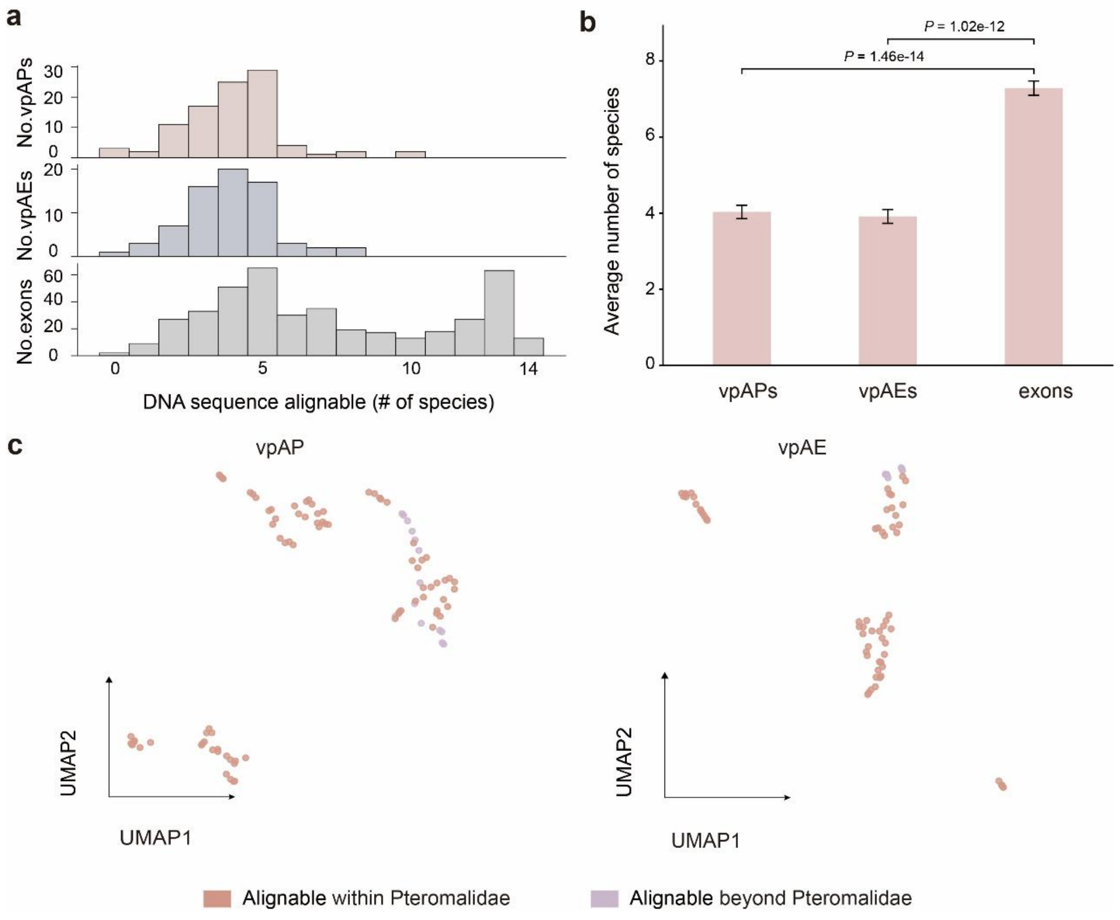


Supplementary Fig. 4. Cross-species alignability of venom-related active promoters, enhancers and venom gene exons. (a) Sequence conservation of vpAPs and vpAEs across parasitoid wasp species, compared to the coding regions of venom genes. (b) Comparison of the average number of alignable species for venom-related active promoters, venom-related active enhancers and exons of venom genes. Statistical significance was calculated using one-sided Mann Whitney U-tests. (c) UMAP visualizations of vpAPs and vpAEs, coloured by the phylogenetic span of sequence alignability, distinguishing elements alignable only within Pteromalidae from those alignable beyond Pteromalidae.

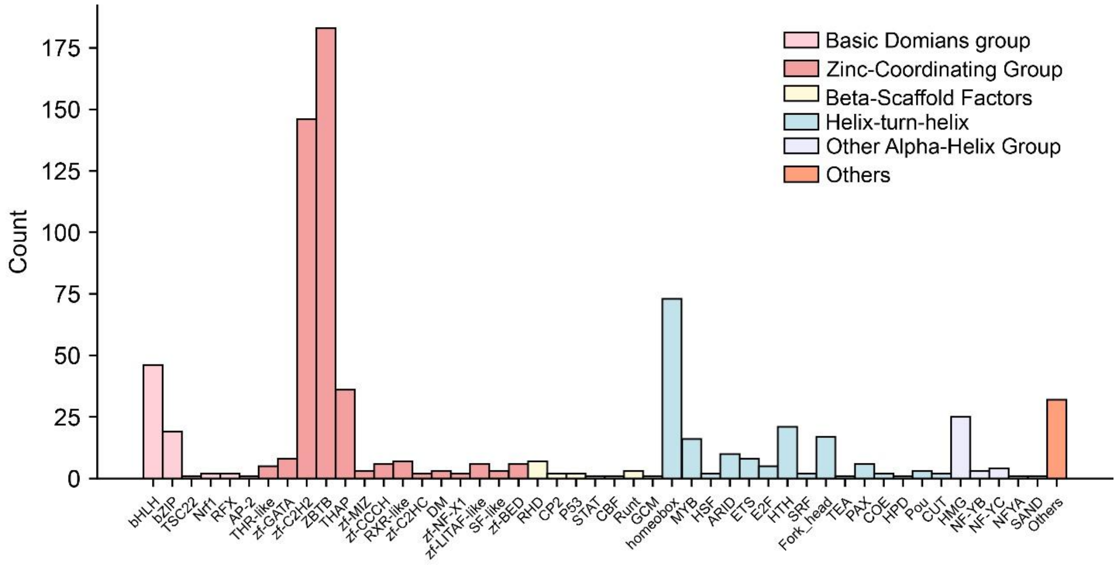


Supplementary Fig. 5. Genome-wide identification and classification of transcription factors in *P. puparum*.

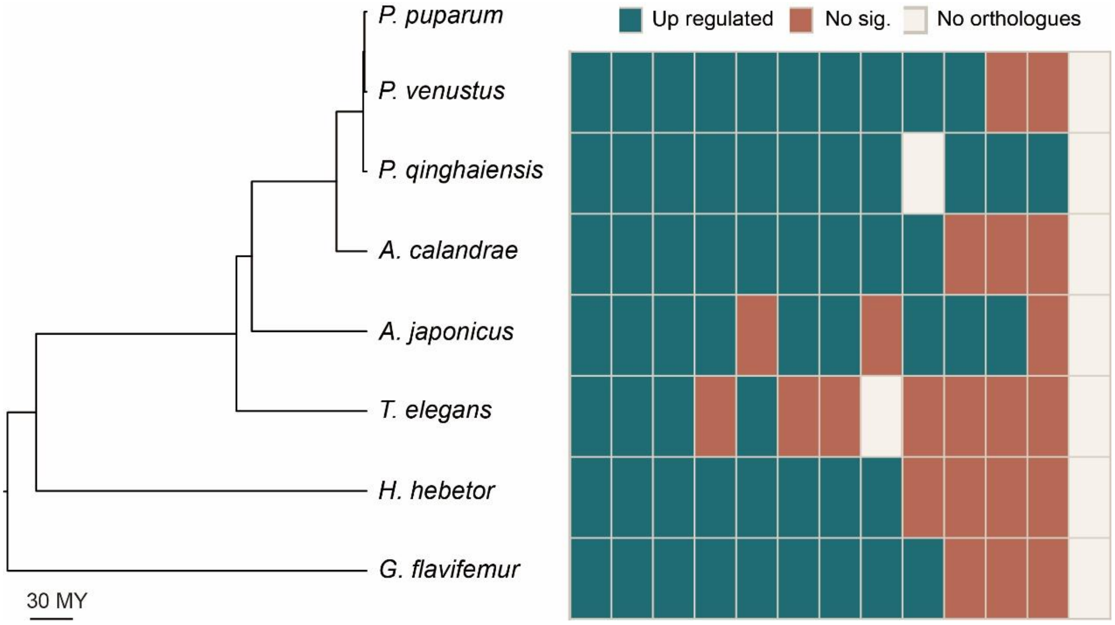


Supplementary Fig. 6. Evolutionary conservation of transcription factors associated with venom gene regulation. The phylogenetic tree was adopted from Supplementary Fig. 4. For each species, differential expression of each TF orthologue was assessed by comparing expression in the venom gland with that in other tissues (i.e. carcass). TFs with log2(fold change) > 1 and adjusted *P* value < 0.05 were defined as significantly upregulated in the venom gland.
